# Mechanisms of PP2A-Ankle2 dependent nuclear reassembly after mitosis

**DOI:** 10.1101/2024.10.07.617051

**Authors:** Jingjing Li, Xinyue Wang, Laia Jordana, Éric Bonneil, Victoria Ginestet, Momina Ahmed, Mohammed Bourouh, Cristina Mirela Pascariu, T. Martin Schmeing, Pierre Thibault, Vincent Archambault

**Affiliations:** Institute for Research in Immunology and Cancer, Université de Montréal; Département de biochimie et médecine moléculaire, Université de Montréal; Département de chimie, Université de Montréal; Department of Biochemistry, McGill University

**Keywords:** Mitosis, Nucleus, Nuclear Envelope, Ankle2, BAF, Lamin, PP2A, *Drosophila*

## Abstract

In animals, mitosis involves the breakdown of the nucleus. The reassembly of a nucleus after mitosis requires the reformation of the nuclear envelope around a single mass of chromosomes. This process requires Ankle2 (also known as LEM4 in humans) which interacts with PP2A and promotes the function of Barrier-to-Autointegration Factor (BAF). Upon dephosphorylation, BAF dimers cross-bridge chromosomes and bind lamins and transmembrane proteins of the reassembling nuclear envelope. How Ankle2 functions in mitosis is incompletely understood. Using a combination of approaches in *Drosophila*, along with structural modeling, we provide several lines of evidence that suggest that Ankle2 is a regulatory subunit of PP2A, explaining how it promotes BAF dephosphorylation. In addition, we discovered that Ankle2 interacts with the endoplasmic reticulum protein Vap33, which is required for Ankle2 localization at the reassembling nuclear envelope during telophase. We identified the interaction sites of PP2A and Vap33 on Ankle2. Through genetic rescue experiments, we show that the Ankle2/PP2A interaction is essential for the function of Ankle2 in nuclear reassembly and that the Ankle2/Vap33 interaction also promotes this process. Our study sheds light on the molecular mechanisms of post-mitotic nuclear reassembly and suggests that the endoplasmic reticulum is not merely a source of membranes in the process, but also provides localized enzymatic activity.

## INTRODUCTION

In animal cells, the disassembly of the nucleus during mitosis allows the segregation of the genetic material via the spindle apparatus, composed of cytoplasmic microtubules and centrosomes. However, this process of open mitosis complicates the transition from mitosis to interphase, as a new nucleus must reassemble around each set of segregated chromosomes. The mechanisms of nuclear reassembly after mitosis remain incompletely understood (Hampoelz and Baumbach, 2023; Kono and Shimi, 2024; Li et al., 2024; Schellhaus et al., 2016; Ungricht and Kutay, 2017).

The nuclear envelope (NE) is a double membrane that is topologically continuous with the endoplasmic reticulum (ER) (Deolal et al., 2024; Ungricht and Kutay, 2017). The NE is associated with multiple structural proteins that connect it to chromatin and cytoskeleton. Lamins form a semi-rigid cage-like network between chromatin and the inner nuclear membrane while the Barrier-to-Autointegration Factor (BAF) interacts with DNA, Lamins, and transmembrane LEM-Domain proteins within the inner nuclear membrane (Sears and Roux, 2020). Nuclear pore complexes composed of multiple nucleoporins ensure transport across the NE and also interact with structural cytoplasmic and nucleoplasmic proteins (Goldberg, 2017; Petrovic et al., 2022; Shevelyov, 2023). In addition, the LINC complex spans the entire NE to connect chromatin to the cytoskeleton and relay mechanotransduction (Hieda, 2019). During mitotic entry, the phosphorylation of many of these proteins disrupts their interactions, resulting in nuclear envelope breakdown (NEB) (Archambault et al., 2022). BAF is phosphorylated by Vaccinia-Related Kinases (VRKs, NHK-1/Ballchen in *Drosophila*) at its N-terminus, which impedes its binding to DNA (Lancaster et al., 2007; Nichols et al., 2006). Similarly, CDK1, PKC and PLK1 phosphorylate Lamins A/C and B, disrupting their polymerization (Liu and Ikegami, 2020; Velez-Aguilera et al., 2020). As a result of NEB, NE membranes retract into the ER during mitosis.

Correct reassembly of the nucleus after mitosis is critical for cell viability and physiology. During this process, segregated chromosomes gather together and ER membranes are recruited around chromosomes to rebuild the NE (Deolal et al., 2024; Schellhaus et al., 2016; Ungricht and Kutay, 2017). Failure to group chromosomes into a single nucleus results in micronuclei that are prone to rupture and DNA damage (Guo et al., 2020). The timely reformation of the nuclear envelope is essential for reestablishing nucleocytoplasmic transport that is needed for the entire gene and protein expression process, including transcription, mRNA export and ribosome biogenesis (Alberts et al., 2015). Protein Phosphatases 1 (PP1) and 2A (PP2A) dephosphorylate several proteins to promote their interactions and reassociation with the nuclear envelope (Archambault et al., 2022; Huguet et al., 2019). How the activities of the enzymes are coordinated in time and space is incompletely understood.

BAF plays a central role in nuclear reassembly. It forms a dimer that binds DNA after anaphase, thereby cross-linking chromosomes into a single cohesive mass (Samwer et al., 2017). BAF also promotes the recruitment of ER membranes through its interactions with LEM-Domain proteins, and aids in the reassembly of the Lamina (Haraguchi et al., 2008; Haraguchi et al., 2001). The dephosphorylation of BAF is required for its recruitment to segregated chromosomes (Archambault et al., 2022; Sears and Roux, 2020). The B55 regulatory subunit of PP2A promotes this regulation in various animals, including *C. elegans*, *H. sapiens* and *D. melanogaster* (Asencio et al., 2012; Mehsen et al., 2018). In addition, the PP2A-interacting, ER-localized protein Ankle2 (also known as LEM4 and LEM-4L in humans and *C. elegans*, respectively) is also required for the recruitment of BAF to segregated chromosomes in the same three organisms (Asencio et al., 2012; Li et al., 2024; Snyers et al., 2018). In humans, Ankle2/LEM4 contains Ankyrin repeats, a LEM domain, a transmembrane domain, and other predicted structured and unstructured regions (Fishburn et al., 2024). The molecular mechanisms by which Ankle2 functions in nuclear reassembly are unclear (Fig 1A). *ANKLE2* function is crucial for the development of the central nervous system as mutations in this gene are associated with microcephaly (Thomas et al., 2022; Yamamoto et al., 2014). Targeting of Ankle2 by the Zika virus also causes microcephaly (Shah et al., 2018). In *Drosophila*, Ankle2 is required for asymmetric divisions in larval neuroblasts (Link et al., 2019). While Ankle2 impacts the asymmetric division protein machinery, it remains unclear whether the loss of its function in nuclear reassembly also contributes to microcephaly.

**Figure 1.**
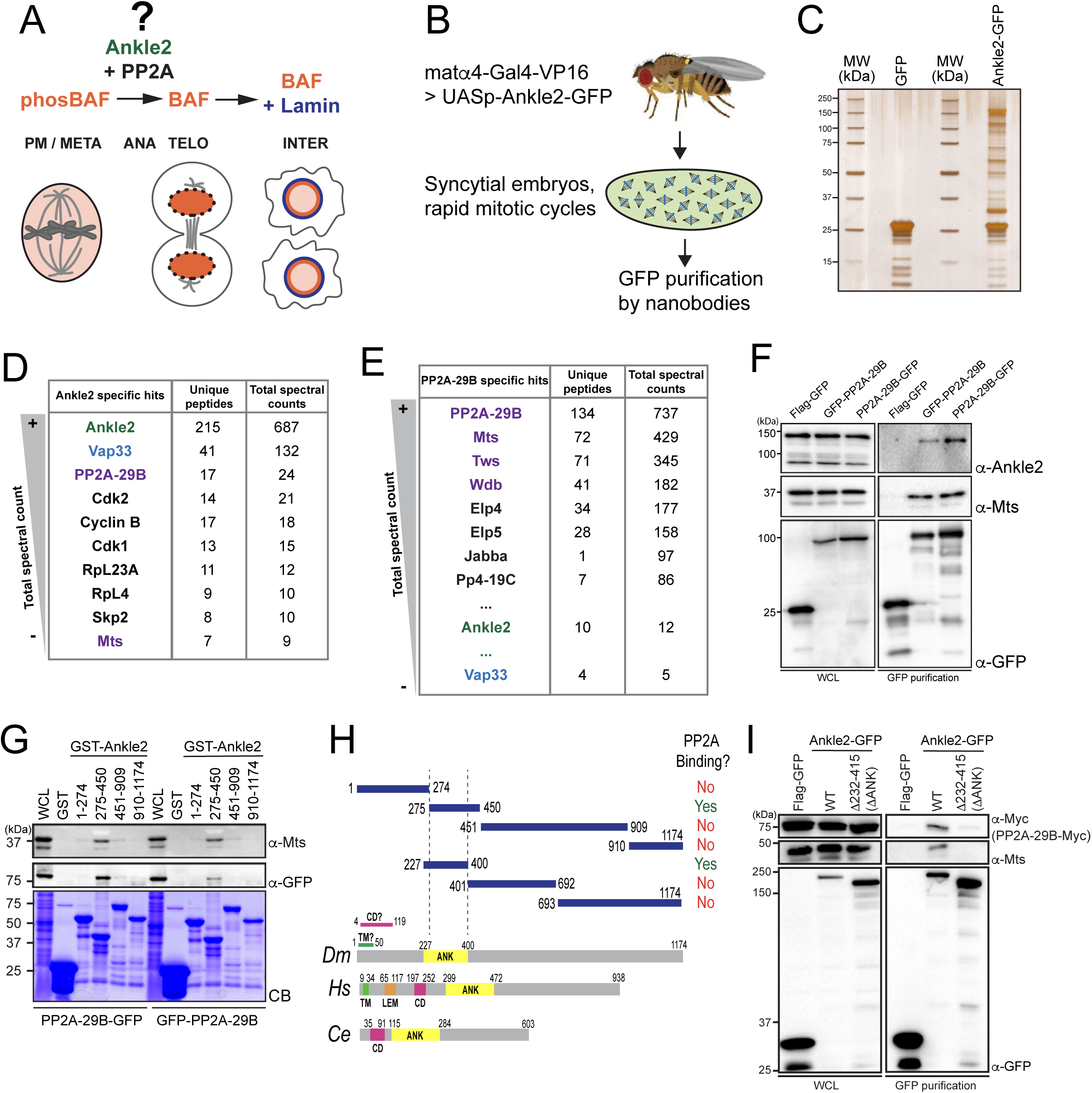
Ankle2 interacts with PP2A through its Ankyrin repeats region. **A.** Ankle2 functions with PP2A to allow the dephosphorylation of BAF and its recruitment to segregated chromosomes. BAF promotes the formation of a single nucleus by cross-bridging chromosomes, and the recruitment of Lamin and ER membranes containing LEM-Domain proteins (not shown). The molecular mechanisms involving Ankle2 in this process are incompletely understood. **B.** Strategy for the identification of Ankle2 interactor proteins *in vivo*. See text for details. **C-E.** Proteins obtained after purification of Ankle2-GFP, PP2A-29B-GFP or GFP from embryos. **C.** Silver-stained gel showing a fraction of the purification products. **D.** Proteins specifically identified with Ankle2-GFP. Proteins with the highest total spectral counts are shown. **E.** Proteins specifically identified with PP2A-29B-GFP after purification from embryos. Proteins with the highest total spectral counts are shown. Ankle2 and Vap33 were also identified as specific interactors further down the list. Purple names: known PP2A subunits. **F.** Ankle2 is specifically co-purified with PP2A-29B. Cells were transfected with the indicated proteins and used in GFP affinity purifications. Products were analyzed by Western blots. WCL: Whole cell extracts. **G.** A region of Ankle2 between amino-acid residues 275-450 is sufficient for interaction with PP2A. GST-fused fragments of Ankle2 produced in bacteria were used in GST-pulldowns with extracts from D-Mel cells expressing PP2A-29B-GFP or GFP-PP2A-29B. Pulled down proteins (PP2A-29B-GFP or GFP-PP2A-29B and Mts) were detected by Western blots. CB: Coomassie Blue. **H.** Top: Summary of results from GST pulldowns testing interactions of Ankle2 fragments with PP2A. Bottom: Primary structures of Ankle2 from *D. melanogaster* (*Dm*), *H. sapiens* (*Hs*) and *C. elegans* (*Ce*). showing known motifs or domains. ANK: Ankyrin domain region; TM: Trans-Membrane motif: LEM: LEM domain. CD: Caulimovirus Domain. **I.** The Ankyrin domain region of Ankle2 between amino-acid residues 232-415 is required for interaction with PP2A. Cells were transfected with the indicated proteins and used in GFP affinity purifications. Products were analyzed by Western blots. WCL: Whole cell extracts.

To investigate Ankle2 molecular functions, we conducted studies in *Drosophila*. We found that Ankle2 forms a complex with the PP2A structural and catalytic subunits, in a mutually exclusive manner with other known regulatory subunits. Our phosphoproteomic analysis validates that BAF dephosphorylation depends on Ankle2. We also identified and characterized a novel interaction of Ankle2 with Vap33, a VAP family protein that promotes Ankle2 localization to the ER during both nuclear reassembly and interphase. We show that the interactions of Ankle2 with PP2A and Vap33 promote BAF recruitment to chromosomes after anaphase. Moreover, the Ankle2/PP2A interaction is essential for post-mitotic nuclear reassembly and development *in vivo*. We propose that Ankle2 functions as a novel PP2A regulatory subunit required for BAF dephosphorylation, and that Vap33-dependent anchoring of Ankle2 to the ER promotes this function in a localized manner during post-mitotic nuclear reassembly.

## RESULTS

### Ankle2 interacts with PP2A through its Ankyrin domain

To investigate the molecular functions of Ankle2, we sought to identify its interaction partners. First, we generated D-Mel (D.mel2) stable cell lines expressing GFP-fused Ankle2 (N or C terminal). We performed GFP affinity purifications on lysate of these cells and identified co-purified proteins by mass spectrometry. Cells expressing Flag-GFP were used as control. We found that PP2A-29B (PP2A-A, structural) and Microtubule Star (Mts, PP2A-C, catalytic) were specifically co-purified with both GFP-Ankle2 and Ankle2-GFP (Fig S1A-B and Table S1). To test if this association occurs *in vivo*, we generated transgenic flies for the expression and purification of Ankle2-GFP in early embryos (Fig 1B). The maternal driver matα4-GAL-VP16 was used to activate the expression of UASp-Ankle2-GFP in late oocytes and syncytial embryos. Embryos expressing GFP alone were used as control. Again, we found PP2A-29B and Mts specifically co-purified with Ankle2-GFP (Fig 1C-D and Table S2). Reciprocally, endogenous Ankle2 was co-purified with PP2A-29B-GFP from cells in culture and embryos, along with the regulatory subunits Tws, Wdb and Wrd (Fig 1E-F, S1C-D and Tables S2, S3). Thus, the interaction of Ankle2 with PP2A is conserved between *Drosophila* and humans (Asencio et al., 2012).

To map the region of Ankle2 that binds PP2A, we used a GST pulldown assay. We produced GST-fused fragments of Ankle2 covering its entire sequence in *E. coli* and tested their ability to pulldown PP2A-29B (with GFP-fused to either end) and Mts from D-Mel cells. We initially found that a fragment comprising amino acid (a.a.) residues 275-450 was sufficient to strongly associate with PP2A-29B and Mts, while fragments 1-274, 451-909 or 910-1174 showed no or very weak association with these proteins (Fig 1G-H). The 275-450 region comprises most of the Ankyrin domain. This domain was recently proposed to span a.a. residues 229-399 (Fishburn et al., 2024), and our inspection of a structural model generated by AlphaFold3 (Abramson et al., 2024) suggested a.a. residues 227-400. We found that the Ankyrin domain (a.a. 227-400 tested) was sufficient for association with PP2A (Fig S1E and 1H). To test if the Ankyrin domain region is required for the interaction of Ankle2 with PP2A, we used a co-purification assay in D-Mel cells. Ankle2-GFP WT or with a deletion of the Ankyrin domain were purified and the presence of co-purified PP2A-29B-Myc and Mts was probed by Western blot. In this assay, we deleted a region comprising the predicted Ankyrin domain plus a few additional downstream residues showing conservation with human Ankle2 and *C. elegans* Lem-4L; (ΔANK: residues 232-415 deleted; S1F). We found that PP2A-29B-Myc and Mts were co-purified with Ankle2^WT^-GFP but not with Ankle2^ΔANK^-GFP (Fig 1I). We conclude that the region of Ankle2 comprising the Ankyrin domain is necessary and sufficient to bind the PP2A-29B/Mts complex in *Drosophila*.

### Ankle2 functions as a PP2A regulatory subunit to promote BAF dephosphorylation

We hypothesized that Ankle2 functions as a regulatory protein required for PP2A to dephosphorylate some of its substrates. The knockdown of LEM-4L or Ankle2 results in the hyperphosphorylation of BAF in its N-terminus in *C. elegans* or *Drosophila* cells, respectively (Asencio et al., 2012; Li et al., 2024). To identify sites that become hyperphosphorylated in the absence of Ankle2 at a proteome-wide level, we used a phosphoproteomic approach (Emond-Fraser et al., 2023). We transfected D-Mel cells with dsRNA against Ankle2 or dsRNA Non-Target (NT). Four days later, cells were lysed, proteins were digested with trypsin and phosphopeptides were enriched, and analyzed by quantitative mass spectrometry. Among hyperphosphorylated peptides upon Ankle2 depletion, we found the expected BAF peptide phosphorylated at Thr4 and Ser5 (Fig 2A). Phosphorylation at the orthologous sites in human BAF was shown to negatively regulate its ability to interact with DNA, and possibly with LEM-Domain proteins (Nichols et al., 2006). Our observation of these Ankle2-dependent sites in BAF validates our approach since BAF is the only reported protein whose phosphorylation state depends on Ankle2 (Asencio et al., 2012; Li et al., 2024). Moreover, BAF could be the only obligatory substrate of Ankle2-dependent dephosphorylation for cell proliferation as lowering the dose of the BAF kinase NHK-1/Ballchen or expression of an unphosphorylatable mutant form of BAF rescues wing development defects caused by the partial depletion of Ankle2 (Li et al., 2024). Nevertheless, we identified several additional hyperphosphorylated sites in other proteins upon Ankle2 knockdown (Fig 2A and Table S5). Although it is likely that some of the hyper-and hypophosphorylated sites observed reflect indirect alterations of cell physiology upon Ankle2 depletion, our results point at several candidate proteins whose dephosphorylation may depend on Ankle2.

**Figure 2.**
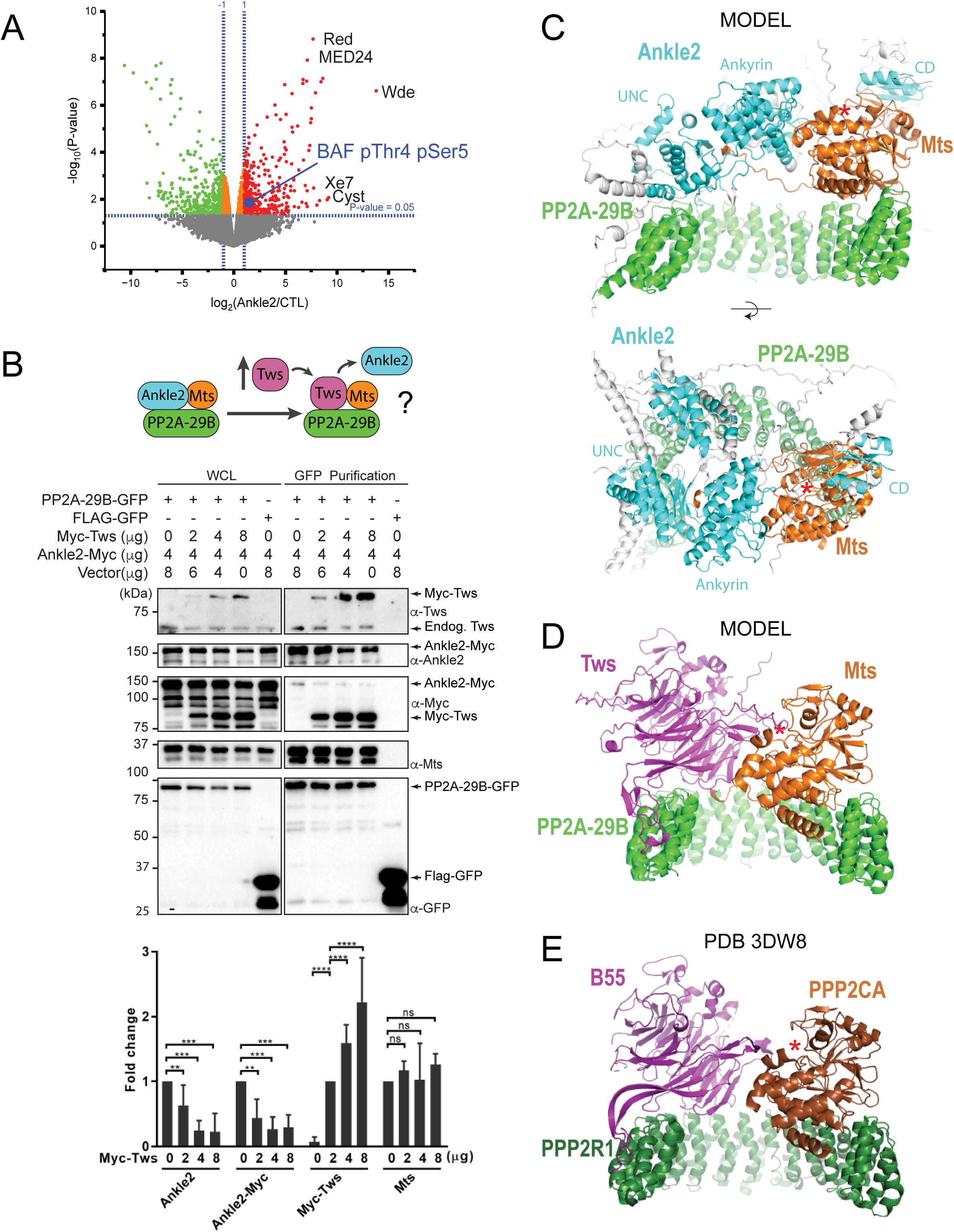
PP2A-Ankle2 promotes BAF dephosphorylation. **A.** Phosphoproteomic analysis identified BAF as being hyperphosphorylated at Thr4 and Ser5 upon Ankle2 depletion. D-Mel cells were transfected with dsRNA against Ankle2 or non-target dsRNA (against bacterial KAN gene). Phosphopeptides from tryptic digests were analyzed quantitatively by mass spectrometry. A few additional significantly hyperphosphorylated proteins are labeled as examples (see Table S5 for the full list). Red, green and orange dots represent peptides with significant (p < 0.05) hyperphosphorylation, hypophosphorylation and little change, respectively. Grey dots indicate peptides with changes below statistical significance. **B.** Competition assay. Top: schematic hypothesis. If Ankle2 occupies the position of a PP2A regulatory subunit, its presence in the PP2A holoenzyme may be outcompeted by Tws (PP2A regulatory subunit). Bottom: the interaction between PP2A-29B-GFP and Ankle2-Myc was monitored by a GFP affinity co-purification assay. Increasing amounts of Myc-Tws plasmid were co-transfected. As a result, increasing amounts of Myc-Tws, and decreasing amounts of Ankle2-Myc, are co-purified with PP2A-29B-GFP. The amounts of co-purified Mts are unchanged. Averages of 3 experiments are shown. All error bars: S.D. * p < 0.05, **p < 0.01, ***<0.001 **** p < 0.0001, ns: non-significant from paired t-tests **C.** AlphaFold3 predicted model of a complex between *Drosophila* Ankle2, PP2A-29B and Mts. Residues of Ankle2 with confidence scores below 0.8 are coloured grey. **D.** Predicted model of a complex between *Drosophila* Tws, PP2A-29B and Mts. **E.** Crystal structure of a complex between human PPP2R1A, PPP2CA and B55 (PDB 3DW8) (Xu et al., 2008). Red asterisks denote the positions of the phosphatase catalytic site.

We hypothesized that Ankle2 functions as a regulatory subunit of PP2A. Interestingly, the structural (PP2A-29B) and catalytic (Mts) subunits of PP2A were detected in complex with Ankle2, while none of the known regulatory subunits of PP2A were detected (Tws, Wdb, Wrd or PR72) (Fig 1D, S1B and Tables S1, S2). To test if Ankle2 interacts with PP2A as a regulatory subunit, we used a competition assay (Fig 2B). We co-transfected cells with Ankle2-Myc and PP2A-29B-GFP or FLAG-GFP. We then proceeded to GFP affinity purifications. As expected, Ankle2-Myc was co-purified specifically with PP2A-29B-GFP. The PP2A regulatory subunit Tws was also co-purified specifically with PP2A-29B-GFP. To test if Tws and Ankle2 are mutually exclusive in the PP2A complex, we co-transfected increasing amounts of Myc-Tws plasmid along with Ankle2-Myc and PP2A-29B-GFP. As a result, increasing amounts of Myc-Tws were co-purified with PP2A-29B-GFP. Conversely, decreasing amounts of Ankle2-Myc were co-purified with PP2A-29B-GFP as the amount of Myc-Tws increased. These results suggest that Tws competes with Ankle2 for binding within the PP2A holoenzyme and reinforce the idea that Ankle2 functions as *bona fide* regulatory subunit of PP2A.

To contextualize and interpret our results, we generated structural predictions of relevant co-complexes using AlphaFold3 (Abramson et al., 2024). In models of a complex of *Drosophila* Ankle2, PP2A-29B and Mts, the Ankyrin domain consistently directly interacts with Mts (Fig 2C shows a representative model), consistent with our pulldown results (Figs 1G and S1E). Interestingly, these models also predict that additional regions of Ankle2 engage in interactions with PP2A. A region previously referred to as the Caulimovirus Domain (CD, Fig 1H) and proposed to mediate an interaction between human Ankle2 and PP2A (Fishburn et al., 2024) is indeed predicted to interact with Mts in our models, on the other side from where the Ankyrin domain binds. Moreover, the modeling shows an adjacent uncharacterized domain (UNC) that is in direct contact with PP2A-29B. However, no interaction with Mts or PP2A-29B was experimentally detected for our GST-fused fragments comprising these regions alone in pulldown assays. It is possible that the UNC and CD domains interact with the PP2A subunits with low affinity. It is also possible that the fragment tested requires additional components to fold into a form that is competent for PP2A binding; in particular, AlphaFold3 predictions show that the complete UNC domain may be made of several segments distal in primary sequence (a.a. residues ∼401-467 + 601-682 + 987-1050). In any case, the deletion of the Ankyrin domain of Ankle2 strongly abrogated its formation of a complex with Mts and PP2A-29B, indicating that it plays a predominant role in the formation of the complex. Importantly, these models display Ankle2 occupying an overlapping position to that of Tws in an analogous PP2A-Tws model (which itself consistent with a crystal structure of human PP2A-B55 (Xu et al., 2008) (Fig 2C-E). These models account for the mutually exclusive binding of Ankle2 or Tws to the PP2A-29B/Mts complex, and are consistent with the □窤沤決已 □□□□□□□□□□□□□□□□□□□□□□□□□℮□□□□□□□□□□□□□□□□Ɋ儀Ɋ帀Ɋ洀

### Ankle2 interacts with Vap33 and they colocalize at the endoplasmic reticulum

Besides PP2A, we discovered that Ankle2 interacts with Vap33, an integral protein of the endoplasmic reticulum (ER) and a member of the VAP family (Kamemura and Chihara, 2019; Murphy and Levine, 2016) (Fig 1D). Mass spectrometry data suggested that Vap33 was co-purified at the highest levels among Ankle2 interactors. Ankle2 was previously reported to localize at the ER and the NE in *Drosophila* (Link et al., 2019). We found that Ankle2-RFP co-localizes with GFP-Vap33 in D-Mel cells (Fig 3A and Video S1). This co-localization is clearest during mitosis, where both proteins are enriched around the presumed spindle in metaphase and around segregated chromosomes in late anaphase/telophase, when NE reassembly takes place. To examine the localization of Ankle2 relative to Vap33 *in vivo*, we generated transgenic flies for the inducible expression of Ankle2-GFP and RFP-Vap33. We expressed the fusion proteins in syncytial embryos and studied their localization. We found that both proteins co-localized throughout the cell cycle (Fig 3B and Video S2). In interphase, they localized to the NE and ER membranes. During mitosis, Ankle2-GFP and RFP-Vap33 became enriched on the spindle envelope until they wrapped around newly forming nuclei in telophase. In addition, both proteins were also enriched at the midbody of the central spindle, consistent with the recruitment of ER membranes at this site (Bobinnec et al., 2003). We were unable to examine the localization of endogenous Ankle2 because the antibodies that we generated failed to reliably detect Ankle2 in immunofluorescence. For the remainder of our study, we used overexpressed Ankle2-GFP, which may not perfectly reflect the localization and function of endogenous Ankle2. However, Ankle2-GFP is functional as it can rescue the phenotypes observed when endogenous Ankle2 is depleted (see below).

**Figure 3.**
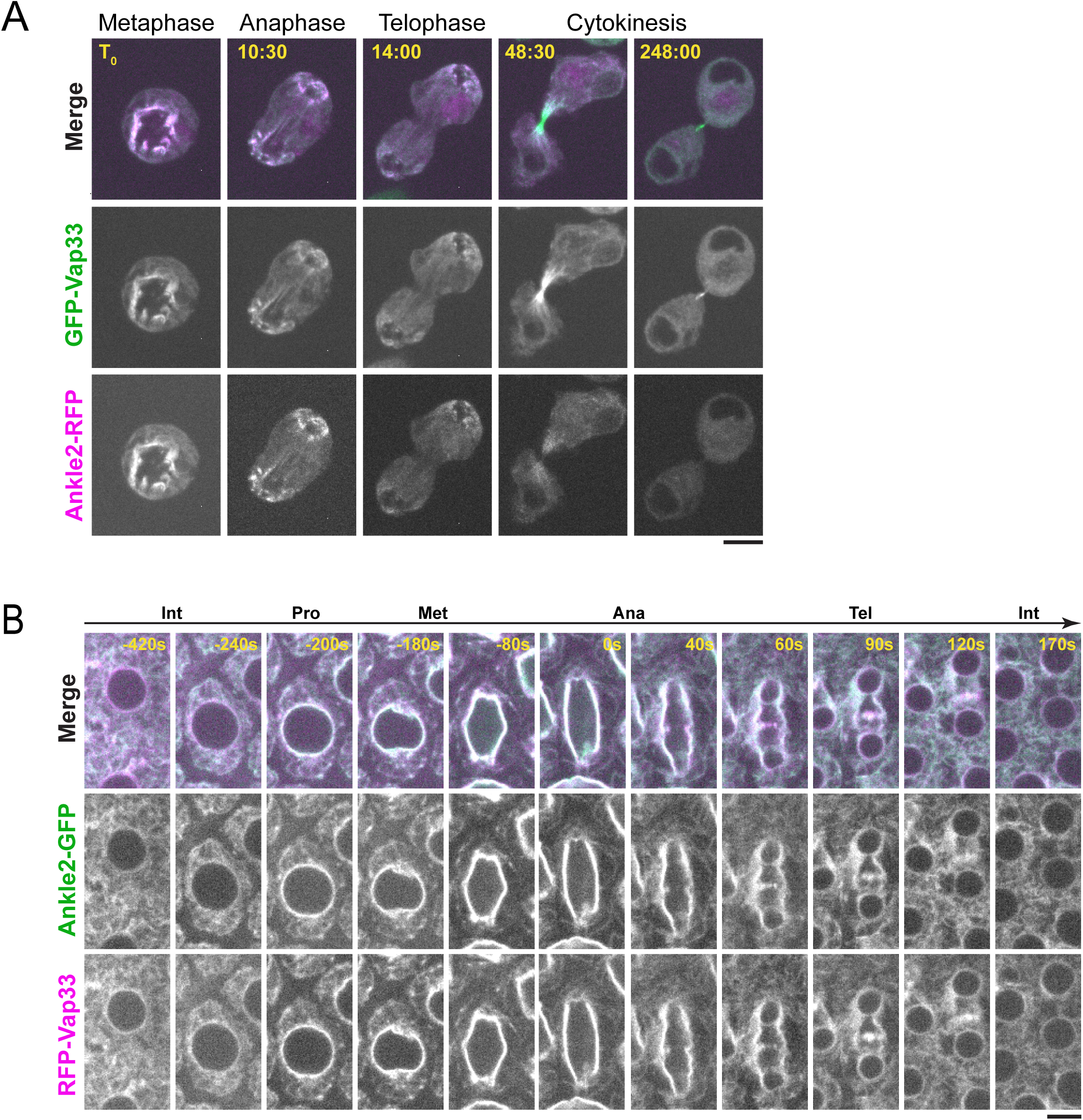
Ankle2 co-localizes with Vap33. **A.** Video images of a D-Mel cell co-expressing Ankle2-RFP and GFP-Vap33 through different stages of cell division. **B.** Video images of a syncytial embryo co-expressing Ankle2-GFP and RFP-Vap33 through different stages of the cell cycle. Scale bars: 5 μm.

### Ankle2 interacts with Vap33 through FFAT motifs

We confirmed the Ankle2/Vap33 interaction in a reciprocal co-purification assay. We found that endogenous Ankle2 was specifically co-purified with GFP-Vap33 or Vap33-GFP from D-Mel cells (Fig 4A). To identify the region of Ankle2 responsible for its interaction with Vap33, we used the GST pulldown described above, probing for the interaction of Vap33-Myc with GST-fused fragments of Ankle2. We found that the C-terminal region of Ankle2 (residues 910-1174) is sufficient to interact with Vap33 (Fig 4B). The VAP family of proteins is anchored to the ER via a transmembrane domain and exposes a globular MSP domain to the cytoplasm (Kamemura and Chihara, 2019; Murphy and Levine, 2016). The MSP domain interacts with proteins that contain FFAT motifs (2 phenylalanines in an acidic tract) (Loewen et al., 2003). To test if Vap33 interacts with Ankle2 in this manner, we introduced mutations in the MSP of Vap33 domain predicted to disrupt its ability to bind FFAT motifs based on results with human VAP (K89D, M91D in Vap33) (Kaiser et al., 2005). We found that this mutant form of Vap33-Myc was unable to interact with GST-Ankle2^910-1174^ (Fig 4C). We then searched for FFAT motifs in Ankle2 using published criteria (Slee and Levine, 2019) and identified three candidate motifs (Fig 4D). One of them was a particularly good fit. Mutation of this FFAT motif (Fm) abolished the interaction of the Ankle2 C-terminal fragment with Vap33 (Fig 4B-C). To test if the interaction occurs in the same way between proteins expressed in *Drosophila* cells, we used a co-purification assay. We transfected different forms of Ankle2-GFP along with Vap33-Myc and proceeded to GFP affinity purifications followed by Western blot for Myc. In this assay, mutation of the first FFAT motif in Ankle2 diminished but did not abolish its interaction with Vap33 (Fig 4E). We therefore tested the contributions of two other additional FFAT-like motifs (FL1 and FL2, Fig 4D). Mutation of FL1 (FL1m) also diminished but did not abolish the interaction of Ankle2 with Vap33, while mutation of FL2 (FL2m) did not perturb the Ankle2-Vap33 interaction. Finally, we found that the combined mutation of the FFAT and FL1 motifs in Ankle2 (Fm+FL1m) abolished its interaction with Vap33. Thus, we conclude that Vap33, via its MSP domain, can interact with Ankle2 through two redundant FFAT motifs. Interestingly, subsequent modeling using AlphaFold3 predicted that Vap33 interacts with Ankle2 preferentially through a contact between the MSP domain of Vap33 and the FL1 motif of Ankle2 (Fig 4F).

**Figure 4.**
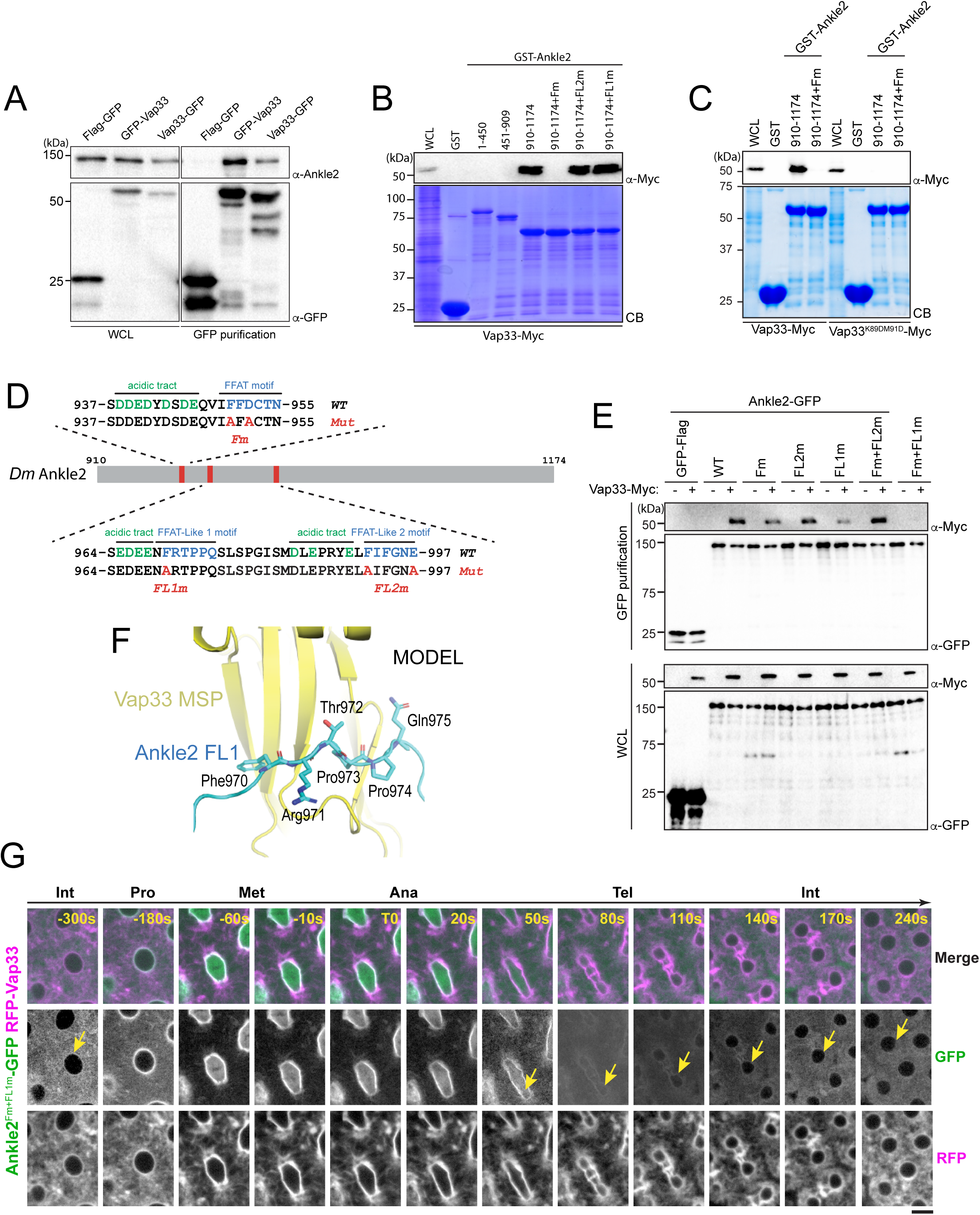
Ankle2 interacts with Vap33 through FFAT motifs. **A.** Cells were transfected with the indicated proteins and used in GFP affinity purifications. Products were analyzed by Western blots. WCL: Whole cell extracts. **B-C.** GST pulldown mapping Ankle2-Vap33 interaction. GST-fused fragments of Ankle2 (WT or with the indicated mutations) produced in bacteria were used in GST-pulldowns with extracts from D-Mel cells expressing Vap33-Myc or Vap33^DD^-Myc. Pulldown products were analyzed by Western blots. CB: Coomassie Blue. WCL: Whole cell extracts. **D.** FFAT and FFAT-Like motifs identified (green) in the C-terminal end of Ankle2. Acidic residues are in blue. The mutations introduced (Fm, FL1m and FL2m) are in red. **E.** The FFAT and FFAT-Like 1 (FL1) motifs contribute to the interaction of Ankle2 with Vap33. Cells were transfected with the indicated proteins and used in GFP affinity purifications. Products were analyzed by Western blots. **F.** AlphaFold3 predicted model of a complex between *Drosophila* Ankle2, Vap33, Mts and PP2A-29B. The image shows a zoomed-in view of the contact between the MSP domain of Vap33 (yellow) and the FL1 motif of Ankle2 (blue). The full heterotetramer structure model is shown in Fig S3B. **G.** The FFAT and FL1 motifs are required for co-localization with Vap33 during telophase and interphase. Video images of a syncytial embryo co-expressing Ankle2^Fm+FLm1^-GFP and RFP-Vap33 through different stages of the cell cycle. To be compared with Figure 3B. Note the delocalization of Ankle2^Fm+FLm1^-GFP at the nuclear envelope indicated by the yellow arrows. Scale bars: 5 μm.

### Vap33 is required for the localization of Ankle2 at the ER and reassembling NE

*Drosophila* Vap33 is an essential protein presumably responsible for interactions of the ER with multiple cytoplasmic proteins and structures (Pennetta et al., 2002). Nevertheless, we attempted to knock down Vap33 by RNAi in the female germline to examine the effects on Ankle2-GFP localization in embryos. Unsurprisingly, we found that those females were sterile and therefore we could not examine embryos. However, we examined ovaries. In control flies, we observed that Ankle2-GFP was weakly localized to nuclear envelopes and strongly localized to the plasma membrane and to ring canals in nurse cells (Fig S2). By contrast, in Vap33 RNAi ovaries, we found that Ankle2-GFP was largely delocalized from these structures. Whether distinct pools of Ankle2 serve specific functions is a question that remains open. Nevertheless, these results suggested that the localization of Ankle2 depends on Vap33.

To test if Ankle2 requires its interaction with Vap33 for its localization, we generated transgenic flies for the expression of Ankle2^Fm+FL1m^-GFP. We then examined its localization in embryos co-expressing RFP-Vap33. In interphase, Ankle2^Fm+FL1m^-GFP failed to localize to the ER and NE like Ankle2-GFP, instead appearing diffuse in the cytoplasm (Fig 4G, to be compared with Fig 3B, and Video S3). Surprisingly, as nuclei entered mitosis, Ankle2^Fm+FL1m^-GFP became strongly localized to the nuclear/spindle envelope, similarly to Ankle2-GFP. However, while Ankle2-GFP was retained at those membranous structures in telophase, Ankle2^Fm+FL1m^-GFP disappeared from them, returning to a diffuse, cytoplasmic localization. We conclude that the interaction of Ankle2 with Vap33 is required for the localization of Ankle2 to the ER in interphase, and for its retention at membranes during NE reassembly in telophase.

Since the ER is the source of the reassembling NE after mitosis, we hypothesized that the interaction of Ankle2 with Vap33 could mediate a localized activity of Ankle2-bound PP2A during nuclear reassembly. In this way, PP2A could dephosphorylate BAF (and potentially other substrates) locally to promote its recruitment near the reforming nuclear envelope (Fig S3A). For this model to be valid, Ankle2 should be able to interact with Vap33 and PP2A simultaneously. We used AlphaFold3 to generate structural models of a complex between Vap33, Ankle2, PP2A-29B and Mts (Fig S3B). The most probable models show that Ankle2 can simultaneously interact with Vap33, PP2A-29B and Mts without steric clash. Our attempts to detect a complex containing Vap33 and PP2A by simple co-IP and Westerns in D-Mel cells gave variable and inconclusive results. However, we detected Vap33 in complex with PP2A-29B-GFP purified from embryos (Fig 1E), despite the fact that PP2A-29B-GFP did not appear clearly enriched at the nuclear/spindle envelope in embryos (Fig S3C). To test if we could promote the formation of the complex, we overexpressed Ankle2-RFP in cells expressing PP2A-29B-GFP and Vap33-Myc. We found that more Vap33-Myc was co-purified with PP2A-29B-GFP under these conditions, compared with cells without Ankle2-RFP overexpression (Fig S3D). This co-purification reflected the specific interaction of Ankle2 with Vap33-Myc because it was abrogated with Vap33^DD^-Myc. Consistent with this result, we found that PP2A-29B-GFP became visibly enriched at the NE upon overexpression of Ankle2-RFP in D-Mel cells (Fig S3E). These results suggest that a Vap33-Ankle2-PP2A complex can mediate the recruitment of a pool of PP2A at the NE.

### Requirements of Ankle2 interactions and motifs for nuclear reassembly

To test the requirements of Ankle2 for its functions in mitosis, we used a rescue assay in D-Mel cells in culture. We depleted endogenous Ankle2 by RNAi and expressed different forms of RNAi-insensitive Ankle2-GFP under the copper-inducible metallothionein promotor (Fig 5A). Depletion of Ankle2 alone resulted in nuclear defects including dispersed or aggregated Lamin, fragmented nuclei, DNA devoid of Lamin and hypercondensed DNA (Fig 5B-C). We previously documented these cellular phenotypes, which we showed to occur as a result of defective BAF recruitment to chromosomes and BAF-dependent nuclear reassembly after mitosis (Li et al., 2024). In addition, Western blotting using Phos-tag revealed that BAF became hyperphosphorylated upon Ankle2 depletion (Li et al., 2024). We verified that expression of Ankle2-GFP rescued the nuclear defects and BAF hyperphosphorylation (Fig 5B-D). When examining cellular phenotypes, we scored only GFP-positive cells. The incomplete restoration of unphosphorylated BAF levels upon expression of Ankle2-GFP (WT and Fm+FL1m) in Ankle2-depleted cells is likely due to the inefficient expression of the GFP-tagged protein in a fraction of the cells.

**Figure 5.**
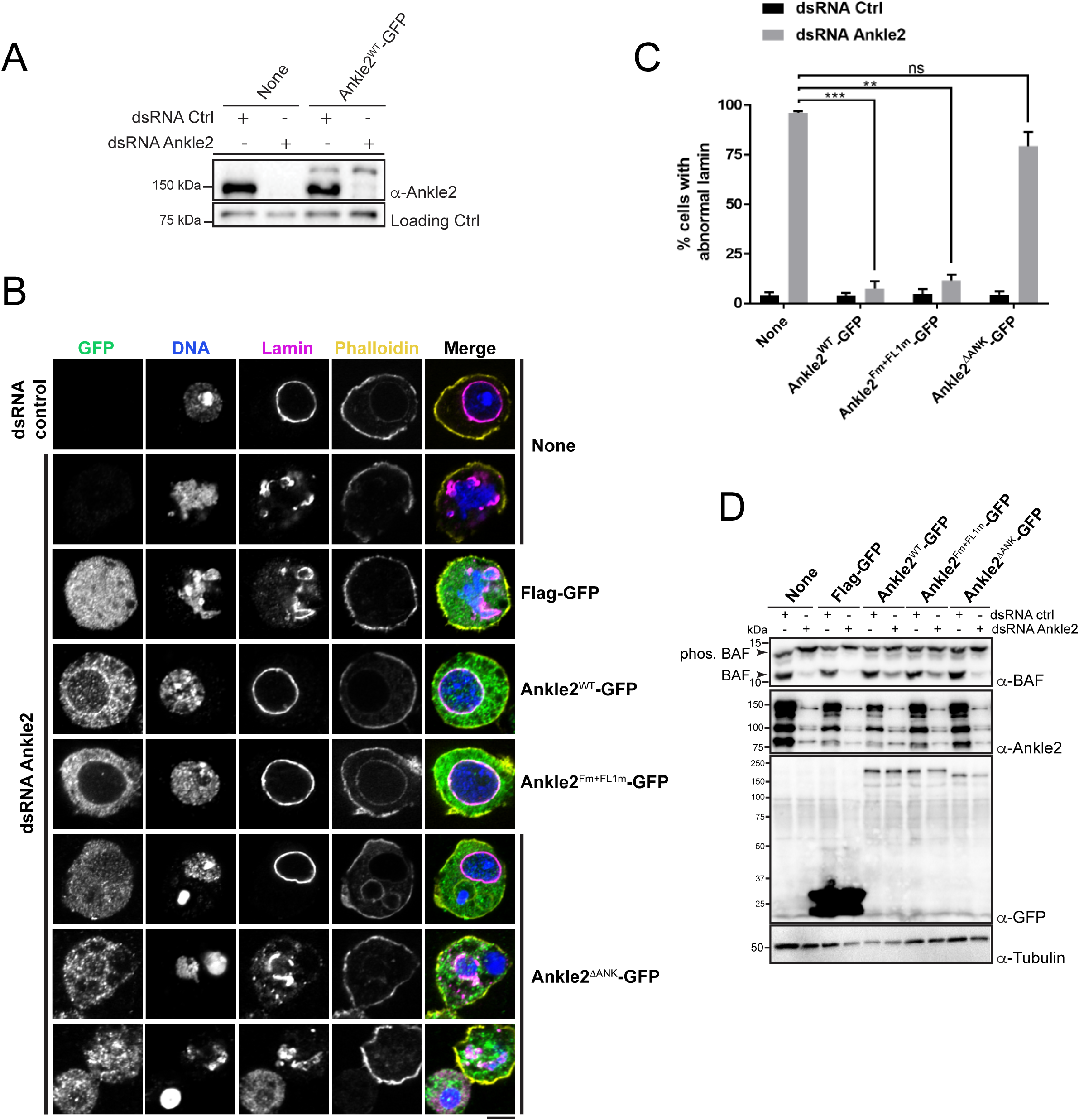
Ankle2 interaction with PP2A but not with Vap33 is required for nuclear reassembly. **A.** RNAi depletion of endogenous Ankle2 and simultaneous expression of RNAi-insensitive Ankle2-GFP. D-Mel cells expressing RNAi-insensitive Ankle2-GFP or not were transfected with dsRNA against Ankle2 or non-target dsRNA (NT, against bacterial KAN gene). Four days later, cells were analyzed by Western blots. Non-specific band is used as loading control. **B.** Immunofluorescence on cells expressing the indicated proteins (right labels) and transfected with the indicated dsRNAs (left labels). GFP fluorescence, DAPI staining for DNA, Lamin (Lamin B) immunostaining and Phalloidin staining for actin are shown. Red frames indicate the presence of nuclear reassembly defects. Scale bars: 5 μm. **C.** Quantification of the abnormal Lamin phenotypes from experiments as in B. Note that Ankle2^WT^-GFP and Ankle2^Fm+FL1m^-GFP, but not Ankle2^ΔANK^-GFP, rescue nuclear defects. Averages of 3 experiments are shown. All error bars: S.D. * p < 0.05, **p < 0.01, ***<0.001 **** p < 0.0001, ns: non-significant from paired t-tests. **D.** Western blot analysis of experiment shown in B-C. The blot at the top was obtained from a gel containing Phos-Tag to increase the resolution between phosphorylated BAF (phos. BAF) and unphosphorylated BAF (BAF). Note that BAF is hyperphosphorylated after Ankle2 depletion and that this is rescued by expression of Ankle2^WT^-GFP and Ankle2^Fm+FL1m^-GFP, but not by Ankle2^ΔANK^-GFP.

To begin to map the essential regions of Ankle2, we tested the ability of Ankle2^1-587^-GFP (N-terminal half) or Ankle2^588-1174^-GFP (C-terminal half) to rescue the nuclear phenotypes. We found that both truncated proteins were unable to rescue nuclear defects and BAF hyperphosphorylation, suggesting that each half of Ankle2 contains at least one essential region for its function (Fig S4). The N-terminal half of Ankle2 contains the Ankyrin domain region necessary and sufficient for Ankle2 interaction with PP2A (Fig 1). We found that Ankle2^ΔANK^-GFP was unable to rescue nuclear defects and BAF hyperphosphorylation (Fig 5B-D). These results indicate that the Ankyrin domain is required for the essential function of Ankle2 in BAF dephosphorylation and nuclear reassembly, and suggest that the ability of Ankle2 to interact with PP2A is necessary for this function.

To map more precisely the C-terminal region(s) of Ankle2 required for its function, we made smaller truncations and deletions in Ankle2-GFP, guided in part by sequence conservation between species. We found that deletion of a.a. residues 607-680, 681-937, 938-997, 992-1040 or 1041-1174 partially or completely abrogated the ability of Ankle2-GFP to rescue nuclear defects and BAF hyperphosphorylation (Figs S4-S5). The 938-997 segment of Ankle2 contains the FFAT motifs required for the interaction with Vap33 (Fig 4). However, we found that the expression of Ankle2^Fm+FL1m^-GFP rescued nuclear defects and BAF hyperphosphorylation (Fig 5B-D). These results suggest that the interaction of Ankle2 with Vap33, while it is required for Ankle2 localization to the reassembling NE in embryos, is dispensable for Ankle2 function in BAF dephosphorylation and nuclear reassembly in D-Mel cells in culture.

Disruption of the interaction with Vap33 prevented the localization of Ankle2 at the ER and the reassembling NE in telophase (Fig 4G); however, it did not abolish the localization of Ankle2 at the nuclear/spindle envelope at early stages of mitosis. We looked for another motif that could contribute to the membrane localization of Ankle2. Human Ankle2 contains a predicted trans-membrane (TM) domain in its N-terminus (Fig 1H) (Asencio et al., 2012). Although online prediction tools failed to predict a TM domain in *Drosophila* Ankle2, we noticed a highly hydrophobic segment in its N-terminus (a.a. 14-31). To test if it contributes to the membrane localization of Ankle2, we combined a deletion of the N-terminal 50 a.a. residues (Fig 1H) with the FFAT mutations. The resulting Ankle2^51-1174&Fm+FL1m^-GFP was recruited to the NE during prophase, similarly to Ankle2-GFP and Ankle2^Fm+FL1m^-GFP (Fig S6). Ankle2^51-1174&Fm+FL1m^-GFP also largely rescued nuclear defects and BAF hyperphosphorylation, although less efficiently than Ankle2-GFP (Fig S4). Among various truncated and deleted forms of Ankle2-GFP, only those that lacked C-terminal a.a. residues 938-1174 (Ankle2-GFP 1-587, 1-692 and 1-937) failed to be recruited to the NE during prophase (Fig S6). This region comprises the 3 candidate FFAT motifs identified. These results suggest that Ankle2 relies on its C-terminus for its NE recruitment during prophase, although the FFAT motifs are not strictly required for this recruitment. We do not know how Ankle2 localizes to the nuclear/spindle envelope independently from Vap33 during early stages of mitosis, or the importance of this localization.

Overall, we conclude that in addition to its N-terminal PP2A-interacting Ankyrin domain, Ankle2 requires the integrity of its C-terminal portion for its essential function in nuclear reassembly. The C-terminus of Ankle2 mediates its interaction with Vap33, which is crucial for the localization of Ankle2 to the reassembling NE in telophase. However, our results suggest that the Ankle2/Vap33 interaction is not strictly required for Ankle2’s function in D-Mel cells.

### Interactions of Ankle2 with PP2A and Vap33 promote BAF recruitment to chromosomes during nuclear reassembly

To test the requirements of Ankle2 interactions *in vivo*, we expressed Ankle2-GFP transgenes in the female germline and early embryos and simultaneously knocked down endogenous Ankle2 using the maternal matα4-GAL-VP16 driver (Fig S7). RFP-BAF was co-expressed. We then imaged mitosis and monitored the dynamics of the fluorescent proteins (Fig 6A top, Video S4). As previously observed, RFP-BAF was enriched at the nuclear/spindle envelope during interphase and early mitosis, likely due to its interactions with LEM-Domain transmembrane proteins (Emond-Fraser et al., 2023). RFP-BAF also largely colocalized with Ankle2-GFP. After anaphase, RFP-BAF transferred to chromatin, becoming visible at the inner core region of the reassembling nuclei, where the chromatin periphery intersects spindle microtubules (Fig 6B), consistent with the role of BAF in nuclear reassembly.

**Figure 6.**
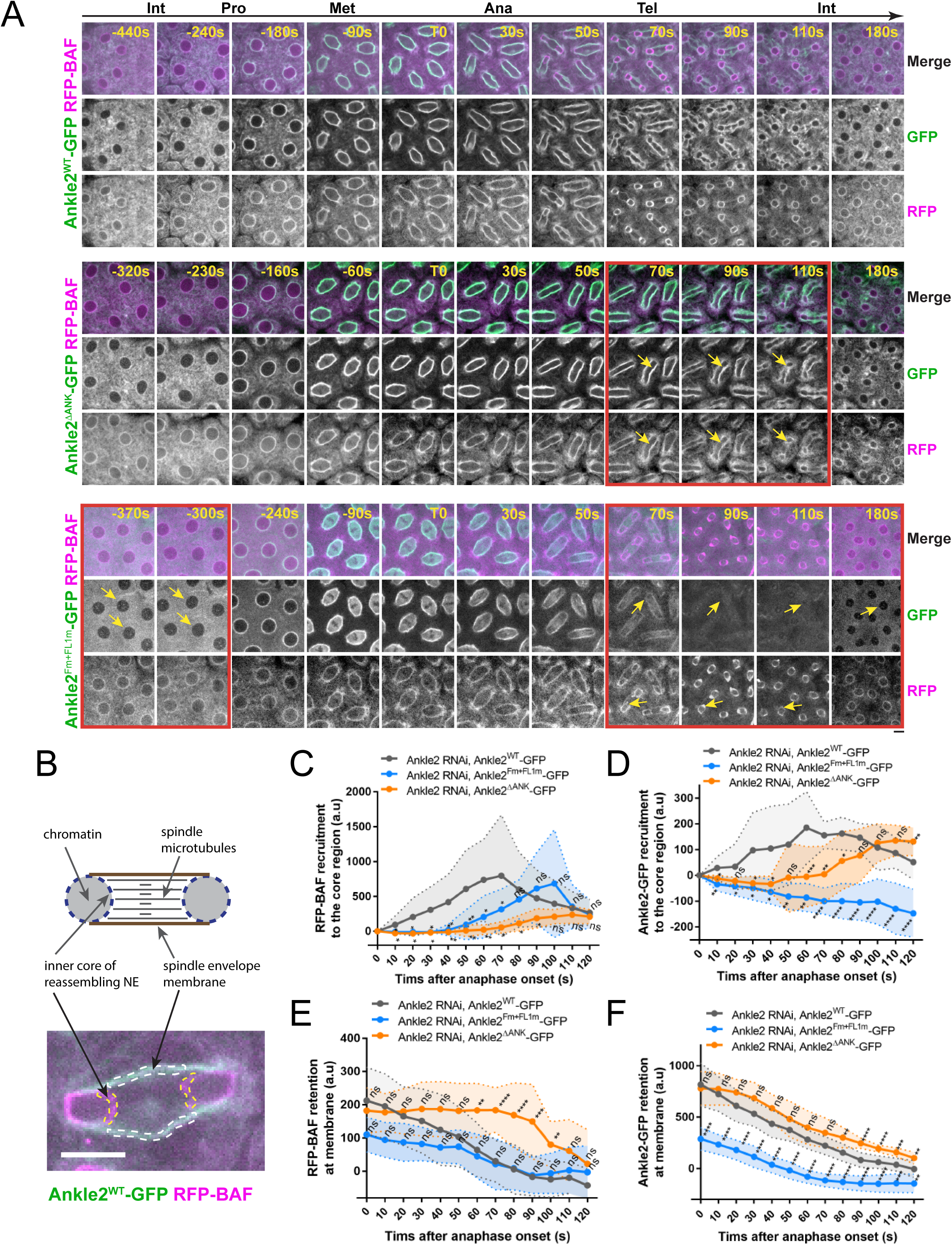
Ankle2 interactions with PP2A and Vap33 promote BAF recruitment at reassembling nuclei. **A.** Syncytial embryos depleted of endogenous Ankle2 and expressing RNAi-insensitive Ankle2^WT^-GFP (top), Ankle2^ΔANK^-GFP (middle) or Ankle2^Fm+FL1m^-GFP (bottom) along with RFP-BAF were imaged through the cell cycle. Times frames where differences are the most pronounced are highlighted by red frames, along with yellow arrows. **B.** Illustration of fluorescence quantifications at specific structures. GFP and RFP fluorescence intensities were quantified at the inner core region of the reassembling nuclei and at the spindle envelope membranes after anaphase. **C-D.** Quantification of the recruitment of RFP-BAF (C) or Ankle2-GFP variants (D) at the inner core region of the reassembling nuclei as a function of time after anaphase onset. **E-F.** Quantification of the retention of RFP-BAF (E) or Ankle2-GFP variants (F) at the lateral spindle envelope membranes as function of time after anaphase onset. Averages of fluorescence intensity from 10 nuclear divisions taken from 5 or 6 embryos are shown per condition. All error bars: S.D. * p < 0.05, **p < 0.01, ***<0.001 **** p < 0.0001, ns: non-significant from unpaired t-tests. Scale bars: 5 μm.

To test the importance of the Ankle2-PP2A interaction, we imaged embryos expressing Ankle2^ΔANK^-GFP. We found that RFP-BAF recruitment to the inner core of reassembling nuclei was strongly diminished and delayed compared to embryos expressing Ankle2^WT^-GFP (Fig 6A middle, Video S5 and Fig 6C). Given that BAF recruitment promotes NE reassembly and Ankle2 is a NE-associated protein, we examined whether the recruitment of Ankle2^ΔANK^-GFP to the inner core region of the reassembling NE was compromised. Fluorescence quantification of NE-associated proteins at the inner core region allows monitoring of NE reassembly, as this region of the reassembling NE is clearly distinct from the spindle envelope in embryos (Emond-Fraser et al., 2023) (Figs 6B and S8A). We found that the recruitment of Ankle2^ΔANK^-GFP to the core region was delayed (Fig 6D), suggesting that the delay in BAF recruitment caused a delay in NE reassembly. Interestingly, RFP-BAF and Ankle2^ΔANK^-GFP were both retained longer at the spindle envelope in embryos expressing Ankle2^ΔANK^-GFP, compared with embryos expressing Ankle2^WT^-GFP (Fig 6E-F). We conclude that the Ankyrin domain, required for the ability of Ankle2 to interact with PP2A, is necessary for the timely recruitment of BAF at reassembling nuclei and ensuing NE reassembly.

To test the importance of the Ankle2-Vap33 interaction, we imaged embryos expressing Ankle2^Fm+FL1m^-GFP (also depleted of endogenous Ankle2) (Fig 6A bottom and Video S6). As before, we observed that Ankle2^Fm+FL1m^-GFP failed to localize to membranes during interphase and telophase, and that it is partially recruited to the nuclear/spindle envelope from prophase to anaphase (Fig S8B). This result rules out the possibility that Ankle2^Fm+FL1m^-GFP localizes to membranes during early mitosis because of a potential complex with endogenous Ankle2. Quantification confirmed that the interaction of Ankle2 with Vap33 is required for the maintenance of Ankle2-GFP at the spindle envelope during telophase and its recruitment to the inner core region of the reassembling NE, at the time when BAF is normally recruited (Fig 6D, F). Moreover, we found that the recruitment of RFP-BAF at reassembling nuclei was delayed in embryos expressing Ankle2^Fm+FL1m^-GFP (Fig 6C). These results suggest that the ability of Ankle2 to interact with Vap33 contributes to the efficient recruitment of BAF at reassembling nuclei.

### The interactions of Ankle2 with PP2A and Vap33 promote its function *in vivo*

We examined the importance of Ankle2 and its interactions for embryonic development. Knockdown of endogenous Ankle2 during late oogenesis using the matα4-GAL-VP16 driver did not significantly diminish the ability of females to lay eggs (Fig 7A). However, none of their embryos hatched, indicating a failure in embryonic development in the absence of maternally contributed Ankle2 (Fig S9A-B). Immunofluorescence revealed that the majority of embryos aborted development in the first mitotic cycle, with a single nucleus/spindle (Fig S9C). FISH for the X chromosome consistently revealed three foci in the polar body and one or two foci in the mitotic spindle, confirming that the spindle observed corresponded to mitosis and not to meiosis or sperm prior to fertilization (Fig S9D-E). Expression of RNAi-insensitive Ankle2-GFP in this background fully rescued embryo hatching. This rescue was strongly abrogated when Ankle2^ΔANK^-GFP was expressed. The weak rescue observed may be in part attributed to the dilution of Gal4 between two UASp elements (UASp-Ankle2 RNAi and UASp-transgene) because a weak rescue was also observed when GFP alone was expressed (Fig 7B). In any case, these results suggest that the interaction of Ankle2 with PP2A, mediated by the Ankyrin domain of Ankle2, is essential during embryo development. Moreover, expression of Ankle2^ΔANK^-GFP in the presence of endogenous Ankle2 did not cause a decrease in embryo hatching, ruling out the possibility of toxicity (Fig 7A). By contrast, expression of Ankle2^Fm+FL1m^-GFP fully rescued embryo hatching, suggesting that the interaction of Ankle2 with Vap33 is dispensable for embryo development (Fig 7B).

**Figure 7.**
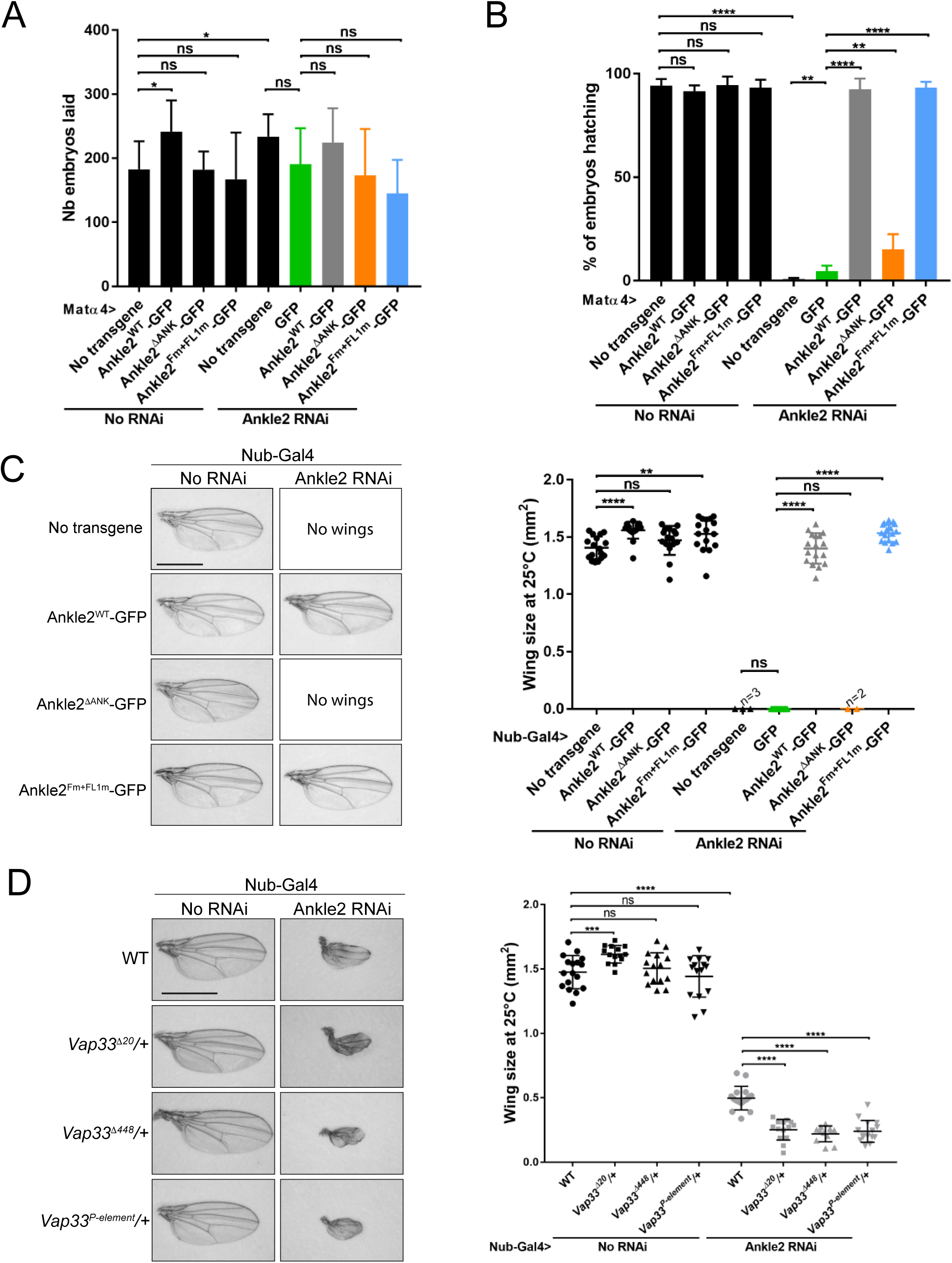
Ankle2 interaction with PP2A but not with Vap33 is required for development. **A.** Number of embryos laid over three days per female of the indicated genotypes. **B.** Percentage of embryos hatching per females of the indicated genotypes. In A-B, the Ankle2 RNAi insertion from line BDSC 77437 was used to deplete endogenous Ankle2 during oogenesis using the maternal matα4-GAL-VP16 driver and RNAi-resistant Ankle2-GFP variants were simultaneously expressed. **C.** Wing development phenotypes from flies of the indicated genotypes. The Ankle2 RNAi insertion from line BDSC 77437 was used to deplete endogenous Ankle2 in the wing pouch using Nub-Gal4 and RNAi-resistant Ankle2-GFP variants were simultaneously expressed at 25°C. Left: Examples of wings. Right: Quantification of wing size. **D.** Mutations in Vap33 enhance a small-wing phenotype caused by Ankle2 depletion. The weaker Ankle2 RNAi insertion from line VDRC100665 was used to deplete endogenous Ankle2 in the wing pouch using Nub-Gal4. Left: Examples of wings. Right: Quantification of wing size at 25°C. Averages of 12-17 wings are shown. All error bars: S.D. * p < 0.05, **p < 0.01, ***<0.001 **** p < 0.0001, ns: non-significant from paired t-tests with Welch’s correction. Scale bars: 1 mm.

To examine the requirements of Ankle2 interactions in a different context, we turned to wing development. Induction of Ankle2 RNAi in the wing disc pouch using Nubbin-Gal4 (Nub-Gal4) resulted in wingless adults. Simultaneous expression of Ankle2-GFP or Ankle2^Fm+FL1m^-GFP rescued the development of adults with wings of the normal size. However, expression of Ankle2^ΔANK^-GFP resulted in no rescue (Fig 7C). These results suggest that the interaction of Ankle2 with PP2A but not with Vap33 is essential for its function during cell proliferation in imaginal wing discs. To test if Vap33 promotes Ankle2 function in a sensitized background, we depleted Ankle2 in the wing pouch using a weaker RNAi line. As a result, smaller adult wings developed (Li et al., 2024). We found that introduction of single mutant alleles of *Vap33* in this background enhances the small wing phenotype. This result suggests that although the Ankle2-Vap33 interaction is not essential, it nevertheless promotes Ankle2 function during wing development (Fig 7D).

## DISCUSSION

Post-mitotic nuclear reassembly is an essential cellular process, yet its molecular underpinnings remain only partially understood. BAF has emerged as a central player required for connections between chromosomes in telophase, for the adhesion of membranes to chromatin through LEM-Domain proteins and for lamina reassembly (Haraguchi et al., 2008; Haraguchi et al., 2001; Li et al., 2024; Samwer et al., 2017). The recruitment of BAF hinges on its dephosphorylation by PP2A, a process dependent on Ankle2 in various animal species including humans, flies and roundworms (Asencio et al., 2012; Li et al., 2024; Mehsen et al., 2018; Snyers et al., 2018). Our study leveraged *Drosophila* as a model to elucidate how Ankle2 functions in this process.

Here we provide several lines of evidence suggesting that Ankle2 functions as a regulatory subunit of PP2A. We found that Ankle2 uses its Ankyrin domain to interact with PP2A. Ankyrin repeats are known to mediate protein interactions (Li et al., 2006). AlphaFold3 predicts an interaction of this domain with PP2A that would position Ankle2 as a regulatory subunit. Indeed, the presence of Ankle2 in the PP2A complex would be mutually exclusive with the presence of known regulatory subunits including B55 (Tws in *Drosophila*) and B56, a hypothesis further supported by our results showing that Tws competes with Ankle2 for binding to PP2A. In addition, Ankle2 co-purified PP2A-29B and Mts without any known PP2A regulatory subunits. Previous work indicated that human Ankle2 uses a similar region to interact with PP2A, comprising part of the Ankyrin domain and a segment immediately preceding it, termed the Caulimovirus Domain (CD, Fig 1H) (Asencio et al., 2012; Fishburn et al., 2024). Although the latter region is poorly conserved in *Drosophila*, our modeling suggests that it may participate in the formation of the PP2A-Ankle2 complex (Fishburn et al., 2024) (Fig 2C). Experimental structural determination of the PP2A-Ankle2 is required to reveal the detailed structure of the complex.

Our unbiased phosphoproteomic analysis confirmed that BAF dephosphorylation depends on Ankle2 in *Drosophila* as in *C. elegans* (Asencio et al., 2012; Li et al., 2024). While this dependence has not been directly tested in human cells, BAF recruitment to segregated chromosomes after anaphase, which is known to require BAF dephosphorylation, requires Ankle2 in HeLa cells (Asencio et al., 2012; Sears and Roux, 2020; Snyers et al., 2018). A LEM domain in human Ankle2 mediates an interaction with BAF (Snyers et al., 2018). By contrast, we did not detect an interaction between *Drosophila* Ankle2 and BAF, consistent with the absence of a LEM domain in the former, as for *C. elegans* (Asencio et al., 2012; Fishburn et al., 2024). Moreover, while human Ankle2 was shown to bind and inhibit the BAF counteracting kinase VRK1 *in vitro* (Asencio et al., 2012), we detected no interaction between Ankle2 and NHK-1/Ballchen (VRK1 ortholog) in *Drosophila*. While a putative interaction between Ankle2 and NHK-1 in *Drosophila* could occur transiently, thereby escaping detection, the simplest interpretation of our results is that the loss of Ankle2 causes BAF hyperphosphorylation by preventing its PP2A-dependent dephosphorylation rather than by preventing inhibition of NHK-1. Our previous genetic analysis in the developing wing suggests that BAF is the most critical substrate of PP2A-Ankle2 during cell proliferation. Indeed, wing developmental defects upon Ankle2 depletion are rescued by lowering the dose of NHK-1 or by expressing an unphosphorylatable mutant form of BAF (Li et al., 2024). Ankle2 may enable PP2A to dephosphorylate BAF through a transient interaction independently from the LEM domain and/or by targeting PP2A to a subcellular location that is favorable to the reaction (discussed below). Interestingly, we identified several additional potential substrates of PP2A-Ankle2 (Fig 2A and Table S5). We were so far unable to obtain sufficient amounts of the PP2A-Ankle2 complex to conduct robust *in vitro* phosphatase assays with phosphorylated BAF or other candidate substrates.

We also discovered a novel interaction between Ankle2 and the ER protein Vap33. We show that FFAT motifs in Ankle2 interact with the MSP domain of Vap33. This interaction is required for the localization of Ankle2 to the reassembling NE in telophase, and to the NE and ER in interphase. The recent identification of putative FFAT motifs in human Ankle2 (Neefjes and Cabukusta, 2021) suggests that the interaction between Ankle2 and VAP family proteins of the ER may be conserved.

These findings led us to propose a model where Vap33 targets the PP2A-Ankle2 holoenzyme to ER membranes, promoting BAF dephosphorylation and recruitment in a localized manner during nuclear reassembly. Supporting this model, we detected a complex comprising Vap33, Ankle2, PP2A-29B and Mts. Our results using cells in culture, embryos and developing wings strongly suggest that the Ankle2-PP2A interaction is essential for BAF dephosphorylation and recruitment to nascent nuclei in telophase, as well as for nuclear reassembly and animal development. However, it remains formally possible that the deletion of Ankyrin repeats used to disrupt the Ankle2-PP2A interaction abrogated another, unknown aspect of Ankle2 function. Similarly, we found that the Ankle2-Vap33 interaction, while being less critical, also promotes BAF recruitment during nuclear reassembly in embryos. Conceptually, the novel mechanism we propose implies that the ER is more than a passive source of membranes in nuclear reassembly; it implicates the ER as a carrier of localized enzymatic activity needed for the process.

Future work should examine the functions of other protein interactions and potential substrates of PP2A-Ankle2 that we identified, which could implicate this holoenzyme in the regulation of various cellular processes. Particularly intriguing are the interactions we identified between Ankle2 and Cyclin-CDK subunits. The possibility that Ankle2, while functioning as a PP2A subunit, may also impact Cyclin-CDK functions warrants further investigation. Conversely, Cyclin-CDKs may also contribute to the regulation of PP2A-Ankle2 in the cell cycle, an area that remains to be explored.

## MATERIALS & METHODS

### Plasmids and mutagenesis

*Drosophila* cells expression vectors were generated using Gateway recombination system (Invitrogen). The cDNA of interest was first cloned into the pDONR221 entry vector and then recombined into the destination vector with C-terminal or N-terminal tag containing copper-inducible or constitutively active promoters, pMT or pAc5, respectively. The RNAi resistant (RNAi res) cDNA for Ankle2 was generated by replacing codons of the original cDNA of the longest form of Ankle2 with synonymous codons. The following expression vectors were generated: pAc5-FLAG-GFP, pAc5-FLAG-Tws, pMT-Ankle2- GFP, pMT-Ankle2^Fm^-GFP, pMT-Ankle2^FL1m^-GFP pMT-Ankle2^FL2m^-GFP, pMT- Ankle2^Fm+FL1m^-GFP, pMT-Ankle2^Fm+FL2m^-GFP, pMT-Ankle2^ΔANK^-GFP, pMT-Ankle2-GFP (RNAi res), pMT-Ankle2^ΔANK^-GFP (RNAi res), pMT-Ankle2^Fm+FL1m^-GFP (RNAi res), pMT- Ankle2^Fm1+FL2m+ΔTM^-GFP (RNAi res), pMT-Ankle2^1-587^-GFP (RNAi res), pMT-Ankle2^588-1174^-GFP (RNAi res), pMT-Ankle2^Δ992-1040^-GFP (RNAi res), pAc5-Ankle2-FLAG (RNAi res), pAc5-Ankle2^F1m+FL2m^-FLAG (RNAi res), pAc5-Ankle2^ΔANK^-FLAG(RNAi res), pAc5- Vap33-Myc, pAc5-Vap33^87D89D^-Myc, pAc5-Vap33-GFP, pAc5-Vap33^87D89D^-GFP, pAc5- PP2A-29B-GFP, pAc5-GFP-PP2A-29B, pMT-RFP-BAF^WT^, pMT-RFP-BAF^3A^. Point mutations and deletions in the pDONR of interest were generated using QuickChange Lightning Site-Directed Mutagenesis Kit (Agilent), as described by the manufacturer’s instructions. GST-Ankle2 (WT, 1-274, 275-450, 451-909, 910-1174, 910-1174+Fm, 910-1174+FL1m, 910-1174+FL2m) expression vectors were constructed into the pGEX4T vector by classic cloning process. pUAS-Ankle2-GFP, pUAS-Ankle2-GFP (RNAi res), pUAS-Ankle2^Fm+FL1m^-GFP (RNAi res), pUAS-Ankle2^ΔANK^-GFP (RNAi res), pUAS-PP2A-29B-GFP, pUAS-RFP-BAF, pUAS-RFP-Vap33 and were generated by cloning PCR amplicons of interest into the pUAS-K10attB vector using restriction enzymes, NotI and BamHI.

### Cell culture, transfections and cell lines

D-Mel cells were cultured in Express Five medium (Invitrogen) supplemented with glutamine, penicillin and streptomycin (Wisent) at 25°C. Transfections with plasmids were performed using X-tremeGENE HP DNA Transfection Reagent (Roche) following the manufacturer’s protocol. Stable cell lines were selected by adding 20 ug/ml blasticidin into the cell culture after transient transfections of plasmids of interest. While inducible pMT-based vectors contain the gene coding for blasticidin resistance, pAc5-based vectors were co-transfected with pCoBlast to confer resistance to blasticidin to the cells. 300μM CuSO_4_ were added into medium to induce expression of the plasmids containing pMT promoter, at least 16 hours before use. For RNA interference, dsRNAs were generated from PCR amplicons using a Ribomax Kit (Promega). RNAi non-target (NT) were generated against the sequence of the procaryotic kanamycin resistance gene. 1×10^6^ cells were plated in 6-well plate and treated with 1ml of medium containing 20 μg of indicated dsRNA for 4 days (rescue experiments). Cells were then harvested and analyzed by immunoblotting, immunofluorescence or live-cell imaging.

### Fly genetics

Fly husbandry was done according to standard procedures. Oregon R was used as the WT strain. Transgenic flies for expression of *pUAS-Ankle2-GFP, pUAS-RFP-Vap33,* and *pUAS-PP2A-29B-GFP* were generated by site-directed insertions of the pUAS-K10attB-based vectors on the second chromosome in the attP40 strain (BestGene Inc, Chino Hills, CA, UAS). Transgenic lines for expression of *pUAS-Ankle2-GFP RNAi res* (WT, Fm+FL1m, ΔANK) were generated by site-directed insertions of the pUAS-K10attB-based vectors on the third chromosome in the attP154 strain (BestGene Inc, Chino Hills, CA, UAS). UAS-Ankle2 RNAi (BDRC 77437) and UAS-GFP (BDRC 4776) used in this study were purchased from Bloomington Drosophila Stock Center.

For the genetic rescue experiment, a combination of UAS-Ankle2-GFP (RNAi res, WT and mutant) and Ankle2 RNAi (BDRC 77437) were generated using a pair of second and third balancer chromosomes. UAS-GFP (BDSC 4776) was used as control for ruling out the possible effect due to Gal4 dilution. Expression of transgenes in the embryo and developing wings was driven by matα4-Gal4-VP16 (BDRC 7062) and Nubbin-Gal4 (BDRC 86108). All crosses were performed at 25°C with 60-70% humidity. Fertility tests were carried out by scoring eggs on the grape juice agar from a single female being crossed with three males for 24 hours.

### Protein purifications from *Drosophila* cells and fly embryos

GFP affinity purifications were performed from *Drosophila* cells and fly embryos, as described (Emond-Fraser et al., 2023). Briefly, cells stably expressing pAc5-FLAG-GFP, pMT-GFP-Ankle2 and pMT-Ankle2-GFP were harvested from four confluent 175cm2 flasks and resuspended in lysis buffer (75 mM K-HEPES pH7.5, 150 mM NaCl, 2 mM EGTA, 2 mM MgCl_2_, 1 mM DTT, 10 µg/ml aprotinin, 10 µg/ml leupeptin, 1 mM PMSF, 5% glycerol, 0.5% Triton X-100). Embryos were collected every 2 hours and dechorionated in 50% bleach, washed in PBS and frozen in liquid nitrogen. Embryos were then crushed in lysis buffer as above. Cell and embryo lysates were incubated for 20 minutes at 4°C on a wheel, centrifuged at max speed for 15 minutes at 4°C, and subsequently incubated with GFP-trap nanobeads (Chromotek) for 2 hours. Beads were firstly washed 5 times with lysis buffer and then washed additionally 5 times in PBS with protease inhibitors (1 mM PMSF, 10 μg/ml aprotinin and 10 μg/ml leupeptin) before sending for mass spectrometry. A small portion of the sample was also eluted in Laemmli buffer and analyzed by SDS-PAGE with silver nitrate staining.

### Mass spectrometry

For proteomic analyses, samples were reconstituted in 50 mM ammonium bicarbonate urea 1M with 10 mM TCEP [Tris(2-carboxyethyl)phosphine hydrochloride; Thermo Fisher Scientific], and vortexed for 1 h at 37°C. Chloroacetamide (Sigma-Aldrich) was added for alkylation to a final concentration of 55 mM. Samples were vortexed for another hour at 37°C. One microgram of trypsin was added, and digestion was performed for 8 h at 37°C. Samples were dried down and solubilized in 5% acetonitrile (ACN)-4% formic acid (FA). The samples were loaded on a 1.5 ul pre-column (Optimize Technologies, Oregon City, OR). Peptides were separated on a home-made reversed-phase column (150-μm i.d. by 200 mm) with a 56-min gradient from 10 to 30% ACN-0.2% FA and a 600-nL/min flow rate on a Easy nLC-1200 connected to a Exploris 480 (Thermo Fisher Scientific, San Jose, CA). Each full MS spectrum acquired at a resolution of 120,000 was followed by tandem-MS (MS-MS) spectra acquisition on the most abundant multiply charged precursor ions for 3s. Tandem-MS experiments were performed using higher energy collision dissociation (HCD) at a collision energy of 34%. The data were processed using PEAKS X Pro (Bioinformatics Solutions, Waterloo, ON) and a Uniprot database. Mass tolerances on precursor and fragment ions were 10 ppm and 0.01 Da, respectively. Fixed modification was carbamidomethyl (C). Variable selected posttranslational modifications were acetylation (N-ter), oxidation (M), deamidation (NQ), phosphorylation (STY). The data were visualized with Scaffold 5.0 (protein threshold, 99%, with at least 2 peptides identified and a false-discovery rate [FDR] of 1% for peptides).

For phosphoproteomic analyses, 500 ug of cell lysate (measured by Bradford assay) were reconstituted in 50 mM ammonium bicarbonate with 10 mM TCEP [Tris(2-carboxyethyl)phosphine hydrochloride; Thermo Fisher Scientific], and vortexed for 1 h at 37°C. Chloroacetamide (Sigma-Aldrich) was added for alkylation to a final concentration of 55 mM. Samples were vortexed for another hour at 37°C. 10 μg of trypsin was added, and digestion was performed for 8 h at 37°C. Samples were dried down in a speed-vac. For the TiO_2_ enrichment procedure, sample loading, washing, and elution were performed by spinning the microcolumn at 8000 rpm at 4 °C in a regular Eppendorf microcentrifuge. The spinning time and speed were adjusted as a function of the elution rate. Phosphoproteome enrichment was performed with TiO_2_ columns from GL Sciences. Digests were dissolved in 400 μL of 250 mM lactic acid (3% TFA/70% ACN) and centrifuged for 5 min at 13000 rpm, and the soluble supernatant was loaded on the TiO2 microcolumn previously equilibrated with 100 μL of 3% TFA/70% ACN. Each microcolumn was washed with 100 μL of lactic acid solution followed by 200 μL of 3% TFA/70% ACN to remove nonspecific binding peptides. Phosphopeptides were eluted with 200 μL of 1% NH4OH pH 10 in water and acidified with 7 μL of TFA. Eluates from TiO_2_ microcolumns were desalted using Oasis HLB cartridges by spinning at 1200 rpm at 4 °C. After conditioning with 1 mL of 100% ACN/0.1% TFA and washing with 0.1% TFA in water, the sample was loaded, washed with 0.1% TFA in water, then eluted with 1 mL of 70% ACN (0.1% TFA) prior to evaporation on a SpeedVac. The extracted peptide samples were dried down and solubilized in 5% ACN-0.2% FA. The samples were loaded on an Optimize Technologies C_4_ precolumn (0.3-mm i.d. by 5 mm) connected directly to the switching valve. They were separated on a home-made reversed-phase column (150-μm i.d. by 150 mm Phenomenex Jupiter C18 stationary phase) with a 120-min gradient from 10 to 30% ACN-0.2% FA and a 600-nL/min flow rate on a Easy nLC-1200 (Thermofisher Scientific, San Jose, CA) connected to an Exploris 480 (Thermo Fisher Scientific, San Jose, CA). Each full MS spectrum was acquired at a resolution of 120,000 and followed by tandem-MS (MS-MS) spectra acquisition for 3 s on the most abundant multiply charged precursor ions. Tandem-MS experiments were performed using higher-energy collisional dissociation (HCD) at a collision energy of 27%. The data were processed using PEAKS X Pro (Bioinformatics Solutions, Waterloo, ON) and a *Drosophila melanogaster* unreviewed Uniprot database. Mass tolerances on precursor and fragment ions were 10 ppm and 0.01 Da, respectively. Variable selected posttranslational modifications were carbamidomethyl (C), oxidation (M), deamidation (NQ), acetyl (N-term) and phosphorylation (STY). The data were visualized with Scaffold 5.0 (protein threshold, 99%, with at least 2 peptides identified and a false-discovery rate [FDR] of 1% for peptides).

### Western blotting and immunofluorescence

Proteins lysates were analyzed on SDS-PAGE and then transferred onto a PVDF membrane. The membrane was then blocked with 5% of milk solution for 1 hour to prevent non-specific binding. Subsequently, the membrane was probed with primary antibodies for 3 hours at 25°C or overnight at 4°C. The following primary antibodies used in this study were: anti-GFP from rabbit (1:5000, Torrey Pine Biolabs), anti-Myc 9E10 from mouse (1:2000, #sc-40, Santa Cruz Biotechnology), anti-FLAG M2 from mouse (1:2000, #F1804, Sigma), anti-tubulin DM1A from mouse (1:5000, #T6199, Sigma), anti-Ankle2 from rabbit (1:1000, custom-made by Thermo Fisher Scientific), anti-Mts from mouse (1:1000, #610555, BD Biosciences), anti-Tws from rabbit (1:2000, custom-made by Thermo Fisher Scientific). After 3 washes of 10 minutes with PBS containing 0,1% Tween (PBST, 0,1%), the membrane was probed with secondary antibody for 30 minutes at room temperature. Peroxidase-conjugated anti-rabbit or anti-mouse from goat (1:5000, Jackson ImmunoResearch) were used as secondary antibodies. The membrane being washed 3 times in PBST, 0,1% was then incubated with Clarity Western ECL Substrate (# 170-5061, Bio-Rad) and subsequently imaged using ChemiDoc^TM^ MP Imaging system. To evaluate the phosphorylation level of the protein of interest, protein lysates were separated by SDS-PAGE in presence of the Phos-tag (Fujifilm WAKO Chemical), following the manufacturer’s instructions. One wash of the acrylamide gel with the transfer buffer containing 1mM EDTA and subsequent second wash with only the transfer buffer were performed before the electrotransfer.

For Immunofluorescence, cells were first fixed with PBS containing 4% formaldehyde for 25 minutes. After three washes of 10min with PBS containing 0,2% Triton X-100(PBST), cells were permeabilized and blocked in PBST containing 1% BSA for 1 hour. Then, cells were incubated with primary antibodies for 2 hours at room temperature, washed three times with PBST, and subsequently incubated with secondary antibodies and DAPI for 1 hour avoiding light. Primary antibodies used in this study were: anti-Lamin from mouse (1:200, DSHB Hybridoma Product ADL84.12), anti-GFP from rabbit (1:1000, Torrey Pine Biolabs). Secondary antibodies from mouse and rabbit were respectively coupled to Alexa-647(1:200, Invitrogen) and Alexa-488 (1:200, Invitrogen).

### Fluorescence *in situ* hybridization (FISH)

*Drosophila* embryos were collected, dechorionated in 50% bleach and washed three times in 0,7% NaCl, 0,05% Triton-100. Eggs were then fixed in methanol: heptane (1:1) and rehydrated successively in methanol: PBS solutions of the ratio of 9:1, 7:3 and 1:1. FISH was performed following procedures as described in, using a probe against the 359-base pair peri-centromeric repeat on the X chromosome. α-Tubulin in the embryos was stained with primary antibodies (YL1/2 from rat at 1:2000, Sigma) and subsequent secondary antibodies (anti-rat Alexa 647 at 1:1000, Invitrogen). DNA was marked with QUANT-IT Oli-green at 1:5000 (#O7582, Invitrogen). Tetrahydronaphthalene was used to mount embryos before being imaged by a confocal microscope.

### GST pulldowns

D-Mel cells expressing Ac5-Vap33-Myc, Ac5-Vap33^87D89D^-Myc, Ac5-PP2A-29B-GFP or Ac5-GFP-PP2A-29B were harvested from a confluent 175 cm^2^ flask. Cells were then lysed in lysis buffer (75 mM K-HEPES pH 7.5, 150 mM KCl, 2 mM EGTA, 2 mM MgCl_2_, 5% glycerol, 0.2% Triton X-100, 1 mM DTT) supplemented with protease inhibitors cocktail as described above at 4°C for 20 minutes. Cell lysates being centrifuged at max speed for 15 minutes were incubated with Sepharose beads bound to GST and GST-Ankle2 (truncated forms) for 1.5 hours at 4°C. Beads were washed 5 times in lysis buffer and subsequently used for SDS-PAGE analysis and Western blot.

### Co-immunoprecipitation (Co-IP)

Pelleted cells from confluent 25 cm^2^ flasks were lysed in 20 mM Tris-HCl pH 7.5, 150 mM NaCl, 2 mM MgCl2, 0.5 mM EDTA, 1 mM DTT, 5% glycerol, 0.5% NP40 Substitute supplemented with protease inhibitors cocktail as described above. Cell lysates being lysed for 15 minutes on a rotating wheel at 4°C were centrifuged at max speed for 10 minutes, and supernatants were incubated with 10μl GFP-Trap agarose beads (Chromotek) for 2 hours at 4°C. Beads were washed four times in 20 mM Tris-HCl pH 7.5, 150 mM NaCl, 2 mM MgCl2, 0.5 mM EDTA, 1 mM DTT, 5% glycerol, 0.1% NP40 Substitute with protease inhibitors. Samples were eluted in Lammeli buffer and analyzed using Western blot.

### Microscopy

A confocal system Leica SP8 was used for imaging of fixed cells and embryos. Acquired images of cells were subsequently processed with the Lightning system. Live imaging of cells and embryos was performed by a spinning disk confocal system (Yokogawa CSU-X1 5000) mounted on a fluorescence microscope (Zeiss Axio Observer.Z1). Cells were cultured on a LabTek II chambered coverglass (#155409, Thermo Fisher Scientific) at least two hours before time-lapse imaging. For live analysis of embryos, 0-2h embryos were dechorionated in 50% bleach and then aligned on a coverslip (#P35G-1.5-14-C, MatTek) and covered with halocarbon oil. Films of embryos were taken every 10s with six confocal sections of about 3μm. To monitor Ankle2-GFP (WT and mutants) and RFP-BAF levels, respectively, the mean intensity of GFP and RFP fluorescence in the region of interest were measured directly with Zen software (Zeiss) at different time points. For each point, the mean intensity of GFP and RFP was normalized by subtraction of mean intensity at anaphase onset (T0) in the core region. Images of adult flies were acquired by stereomicroscope with a camera (Canon). Developing wings of adult flies were dissected and quantified using Fiji software (National Institute of Health).

### Structure Predictions

Prediction of co-complexes was performed using the AlphaFold3 (Abramson et al., 2024) AlphaFold Server (Beta) (https://alphafoldserver.com), with default seed auto-generation and with seed provided, yielding sets of 5 models. Co-complexes modelled are of Mts (NCBI accession NP_001285724.1), PP2A-29B (NP_001027225.1) and Ankle2 (NP_573221.2); Mts, PP2A-29B and Tws (NP_001287269.1); and Mts, PP2A-29B, Ankle2 and Vap33 (NP_570087.1). Each set of 5 models showed self-consistent interactions and a typical example is shown in the figures.

## Statistical analysis

All the graphs and statistical analysis in this study were done by GraphPad software. In all figures, the results of quantifications are expressed as mean ± SD with indicated statistics in the legend. Overall, p values are represented as follows: *p < 0.05, **p < 0.01, ***p < 0.001, ****p ≤ 0.0001, and n.s. (not significant) is p > 0.05.

## Supplemental material

Supplemental materials comprise 9 figures, 5 tables and 6 videos.

## ACKNOWLEDGEMENTS

This work was funded by a Project Grant from the Canadian Institutes of Health Research (CIHR) to VA (175132). JL received a studentship from the *Fonds de Recherche du Québec – Santé* (FRQS). TMS is a member of the *Centre de recherche en biologie structurale*, funded by FRQS Research Centres Grant #288558. We thank Christian Charbonneau for his precious help with the microscopy.

---

**Figure S1.**
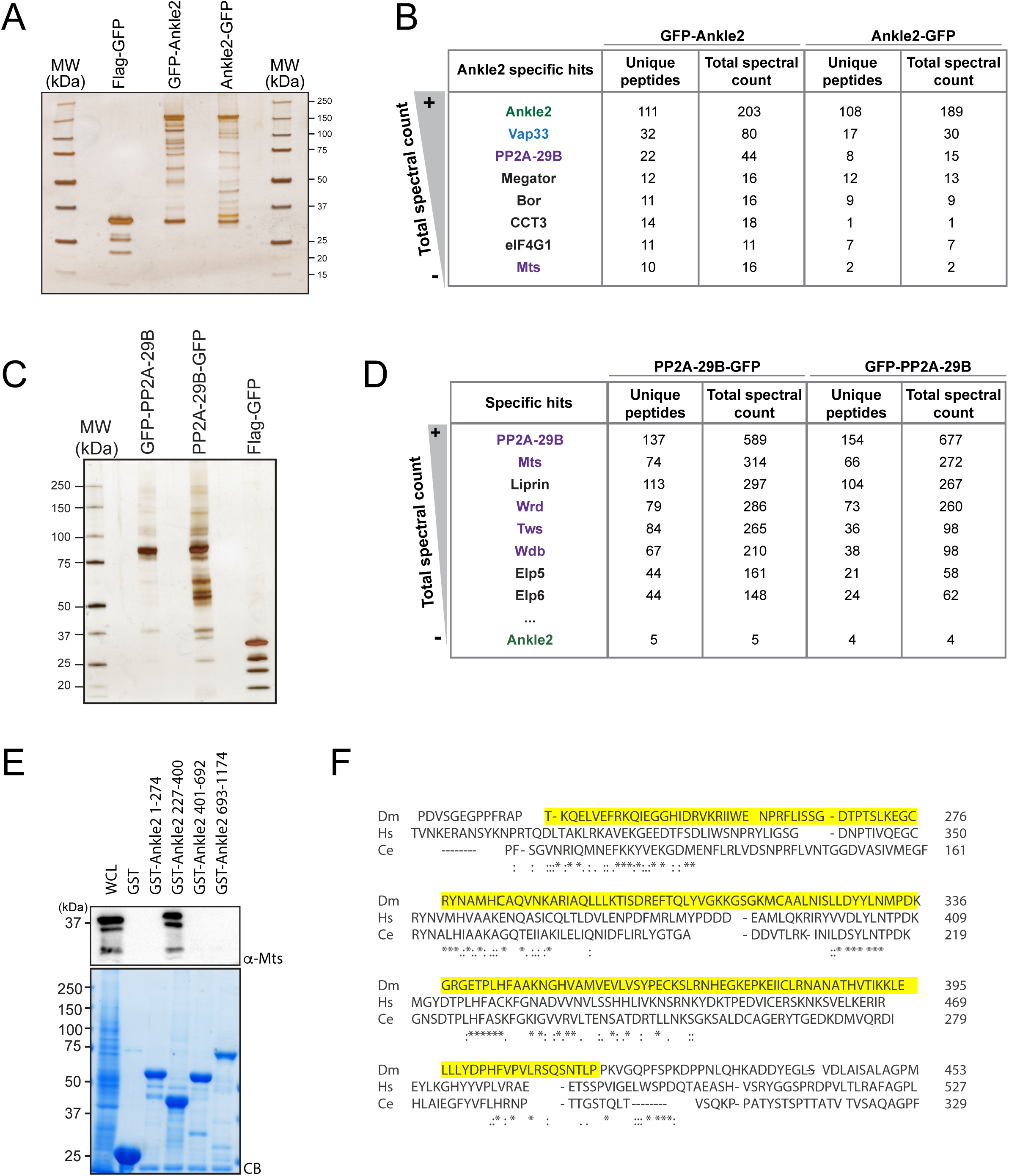
Ankle2 interacts with PP2A through its Ankyrin repeats region - Complement to Figure 1.A-B. Proteins obtained after purification of Ankle2-GFP, GFP- Ankle2 or Flag-GFP from D-Mel cells. **A.** Silver-stained gel showing a fraction of the purification products. **B.** Proteins specifically identified with GFP-Ankle2 or Ankle2-GFP. Proteins with the highest total spectral counts for GFP-Ankle2 are shown. **C-D.** Proteins obtained after purification of PP2A-29B-GFP, GFP-PP2A-29B or Flag-GFP from D-Mel cells. **C.** Silver-stained gel showing a fraction of the purification products. **D.** Proteins specifically identified with PP2A-29B-GFP and GFP-PP2A-29B. Proteins with the highest total spectral counts for PP2A-29B-GFP are shown. Ankle2 was also identified as a specific interactor further down the list. Purple names: known PP2A subunits. **E.** A region of Ankle2 comprising amino-acid residues 227-400 is sufficient for interaction with PP2A. GST-fused fragments of Ankle2 produced in bacteria were used in GST-pulldowns with extracts from D-Mel cells. Mts was detected by Western blot. CB: Coomassie Blue. **F.** Alignment of the Ankyrin domain region of Ankle2 from *D. melanogaster*, *H. sapiens* and *C. elegans*. The alignment was done with full-length sequences using Clustal Omega. The Ankyrin domain region of Ankle2 (a.a. 232-415) deleted in Ankle2^ΔANK^ is shown in yellow.

**Figure S2.**
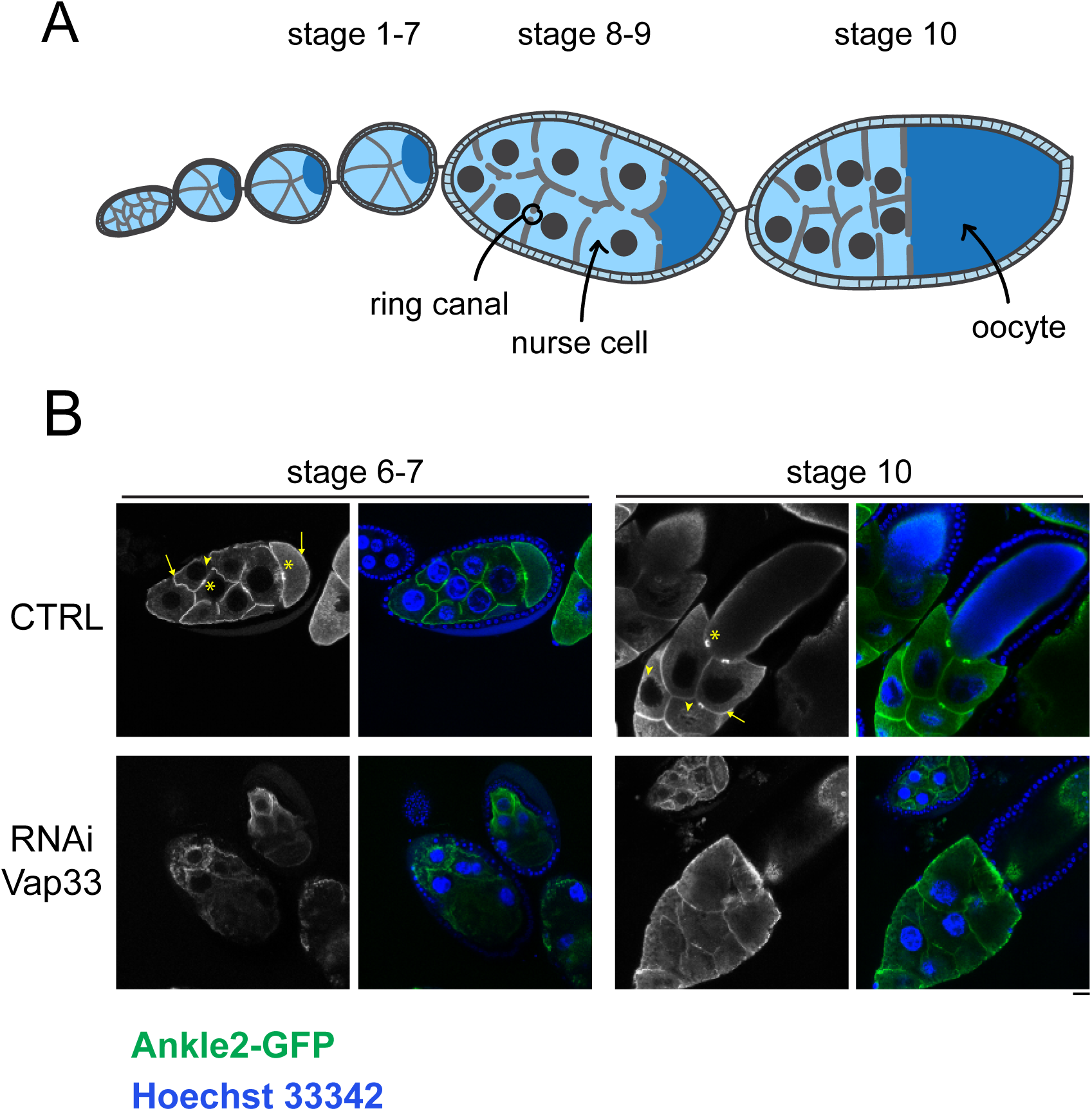
The localization of Ankle2 depends on Vap33 in egg chambers. **A.** Schematic representation of different stages of egg chamber maturation. **B.** Egg chambers expressing Ankle2-GFP from the matα4-GAL-VP16 driver, with or without expression of Vap33 RNAi from the same driver, were stained for DNA with Hoechst 33342. Note that Ankle2-GFP localization around nuclei (arrowheads), the plasma membrane (arrows) and ring canals (asterisks) is abrogated upon Vap33 RNAi. Scale bar: 20 μm.

**Figure S3.**
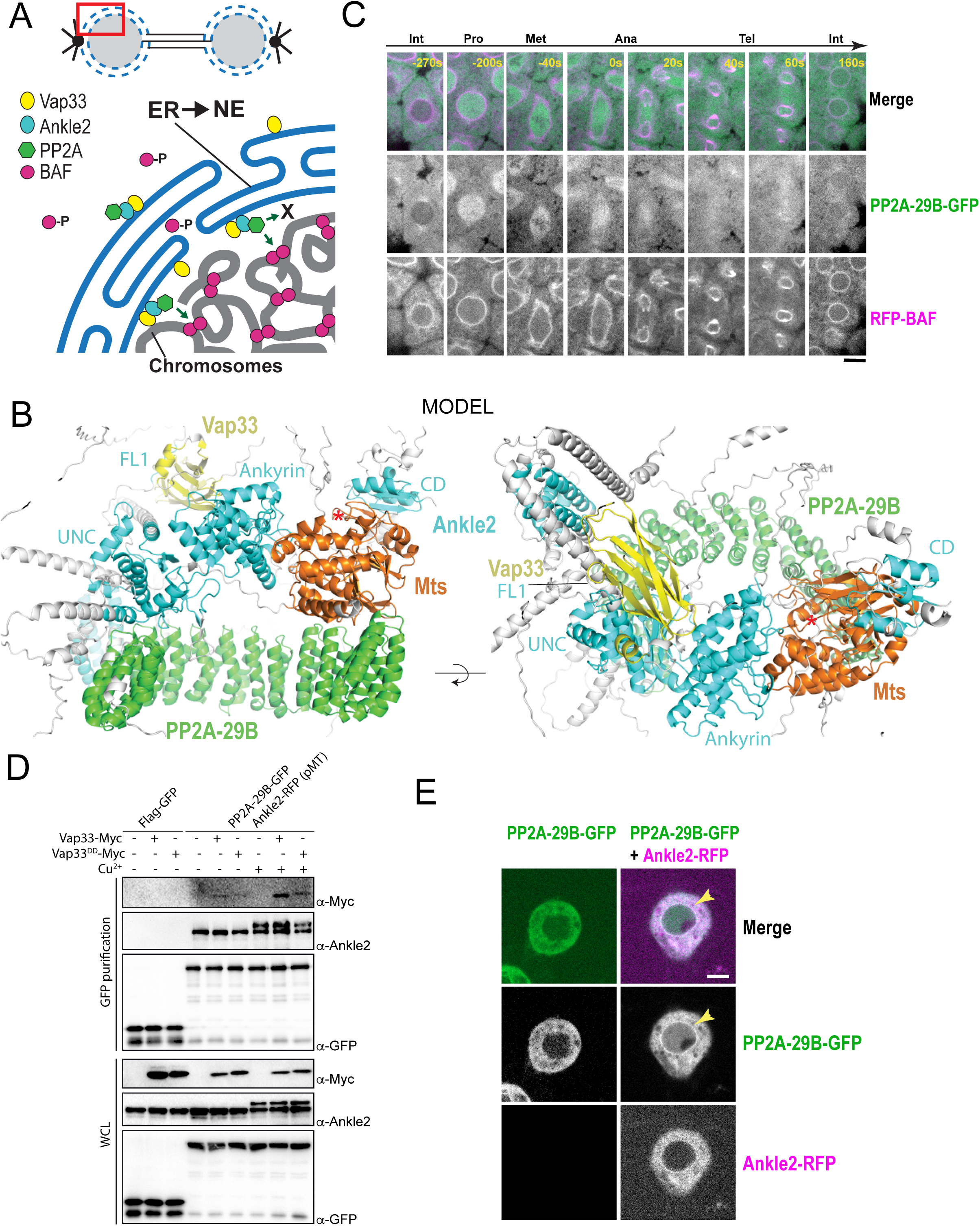
A Vap33-Ankle2-PP2A complex may promote nuclear reassembly in a localized manner. **A.** Hypothetical model discussed in the text. ER: Endoplasmic Reticulum; NE: Nuclear Envelope. **B.** AlphaFold3 predicted model of a complex between *Drosophila* Ankle2, PP2A-29B, Mts and Vap33. Residues with Cα confidence scores below 0.75 are coloured grey. Red asterisks denote the positions of the phosphatase catalytic site. **C.** Imaging of a mitotic cycle in an embryo expressing PP2A-29B-GFP and RFP-BAF. Scale bar: 5 μm. **D.** Overexpression of Ankle2-RFP enhances complex formation between Vap33-Myc and PP2A-29B-GFP. Cells were transfected to express the indicated proteins and submitted to GFP affinity purifications. Cu^2+^ was added to induce the expression of Ankle2-RFP (under the pMT promoter). Products were analyzed by Western blotting. WCL: Whole cell extracts. **E**. Overexpression of Ankle2- RFP enhances the localization of PP2A-29B-GFP to the nuclear envelope (arrows). Scale bars: 5 μm.

**Figure S4.**
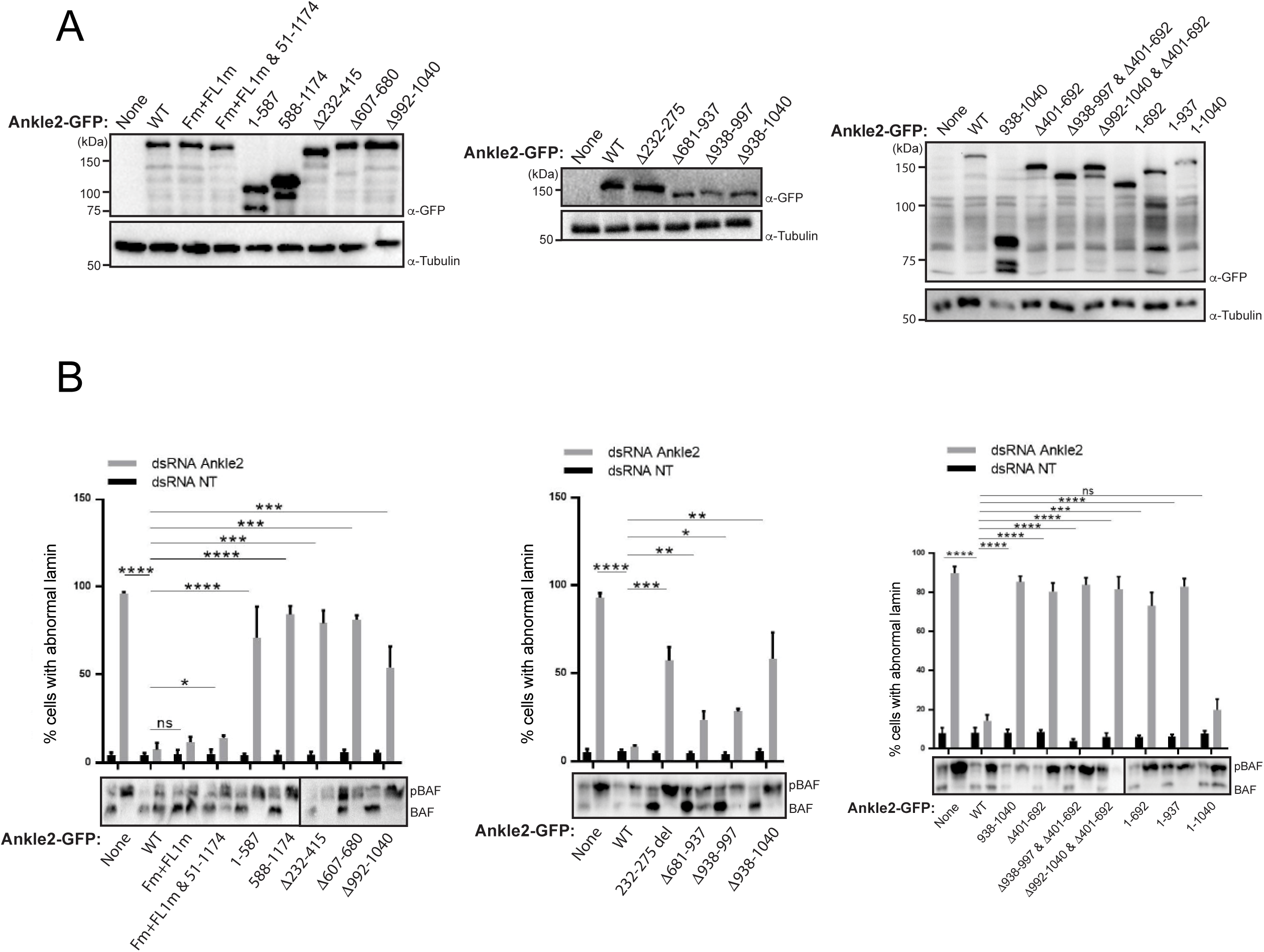
Results of rescue experiments in D-Mel cells in culture. Endogenous Ankle2 was depleted by RNAi and RNAi-insensitive Ankle2-GFP variants were simultaneously expressed as in Figure 6. D-Mel cells expressing RNAi-insensitive Ankle2-GFP variants or not were transfected with dsRNA against Ankle2 or non-target dsRNA (NT, against bacterial KAN gene). **A.** Western blot analyses showing expression of the different variants of Ankle2-GFP. α-Tubulin: loading control. **B.** Quantification of the abnormal Lamin phenotypes from rescue experiments (Top). Western blots (Bottom) were obtained from a gel containing Phos-Tag to increase the resolution between phosphorylated BAF (phos. BAF) and unphosphorylated BAF (BAF). Averages of 3 experiments are shown. All error bars: S.D. * p < 0.05, **p < 0.01, ***<0.001 **** p < 0.0001, ns: non-significant from paired t-tests.

**Figure S5.**
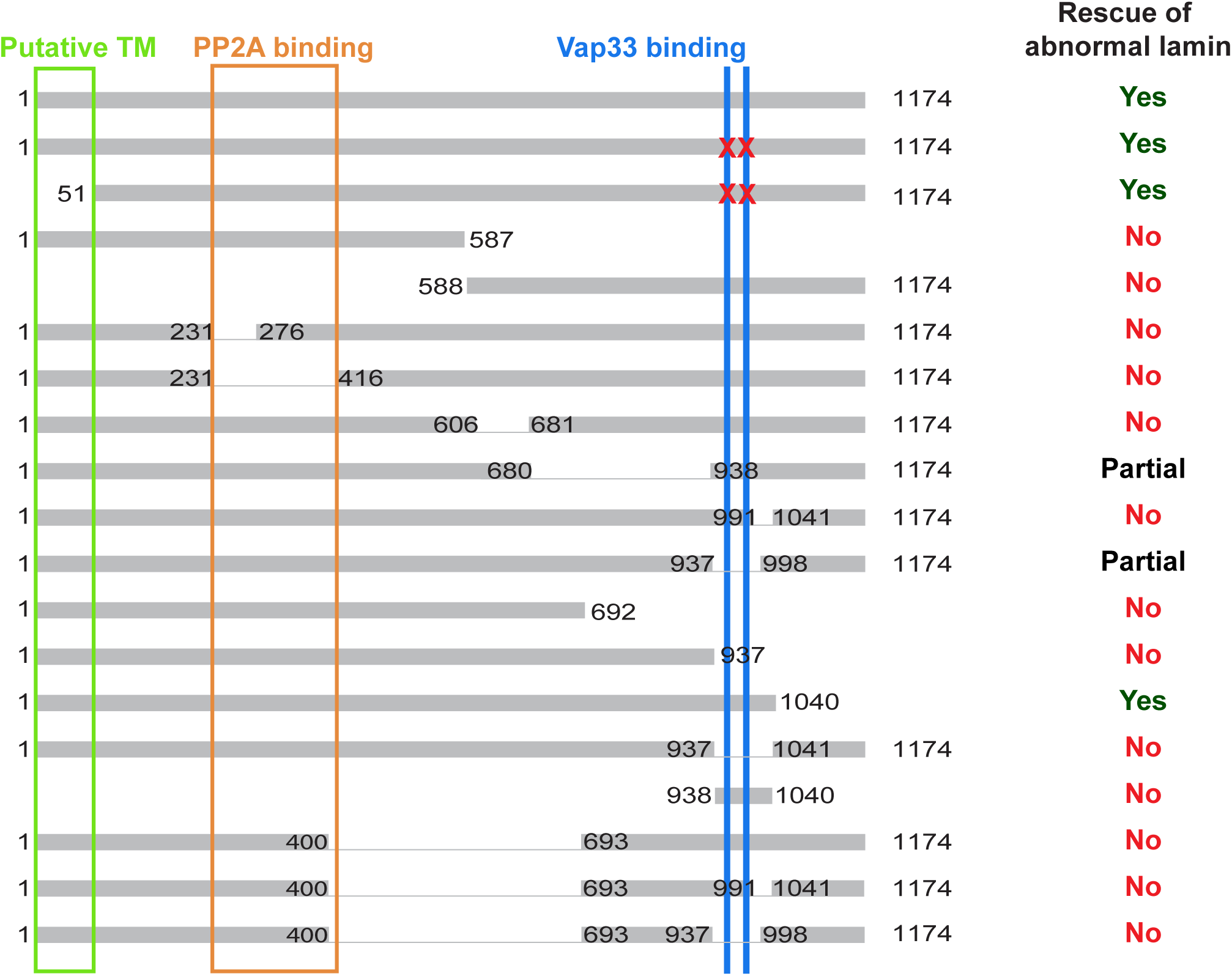
Summary of results for various deletion mutants of Ankle2-GFP assessed for their ability to rescue abnormal Lamin phenotypes. Regions of interest discussed in the text are indicated with colored boxes. Red Xs: mutations of FFAT and FFAT-Like 1 Vap33 binding motifs.

**Figure S6.**
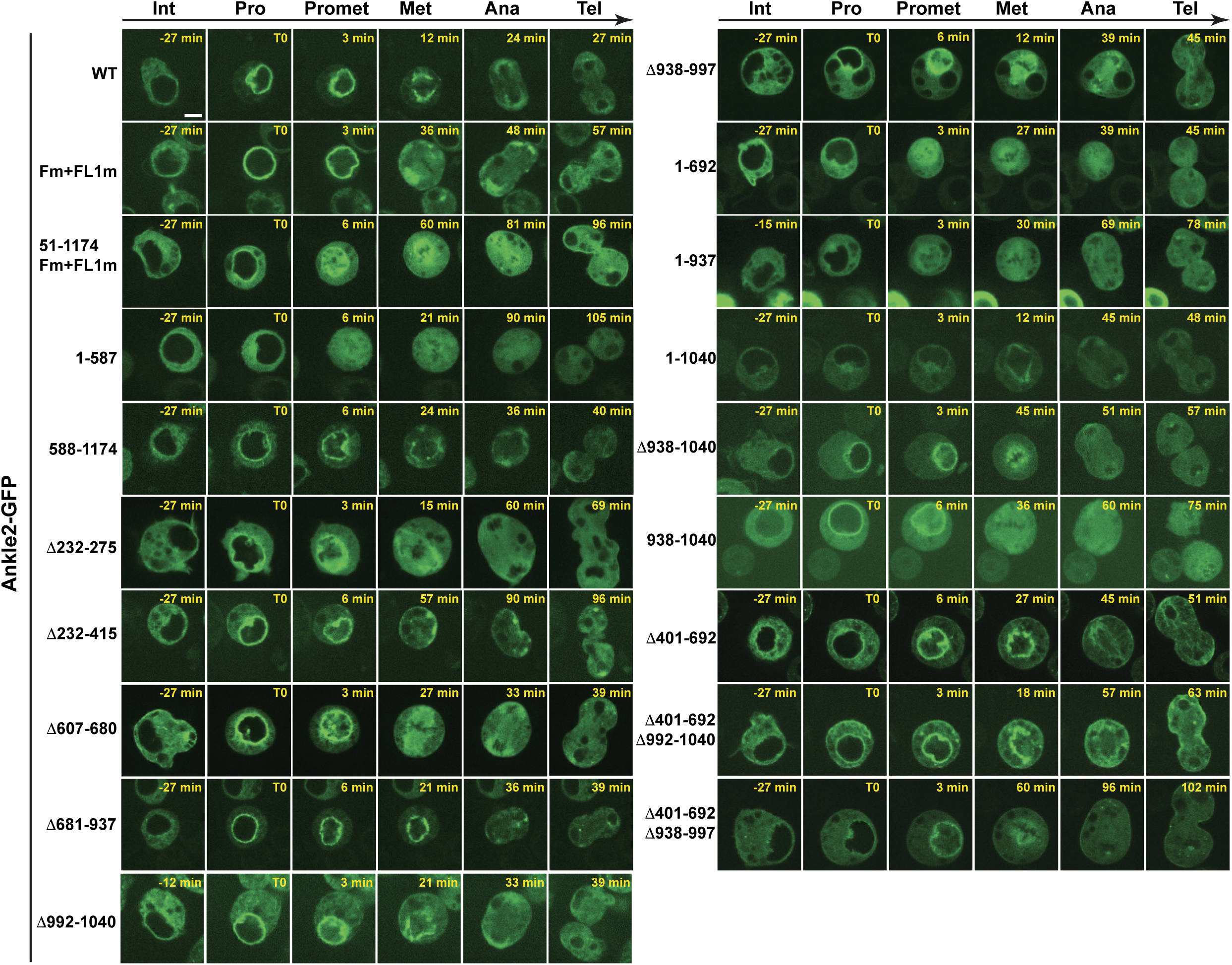
Localization of Ankle2-GFP variants during the cell cycle in D-Mel cells. Cells were imaged on a confocal microscope. Representative images of different stages of the mitotic cycle are shown. Int: Interphase; Pro: Prophase; Promet: Prometaphase; Met: Metaphase; Ana: Anaphase; Tel: Telophase. Scale bar: 5 μm.

**Figure S7.**
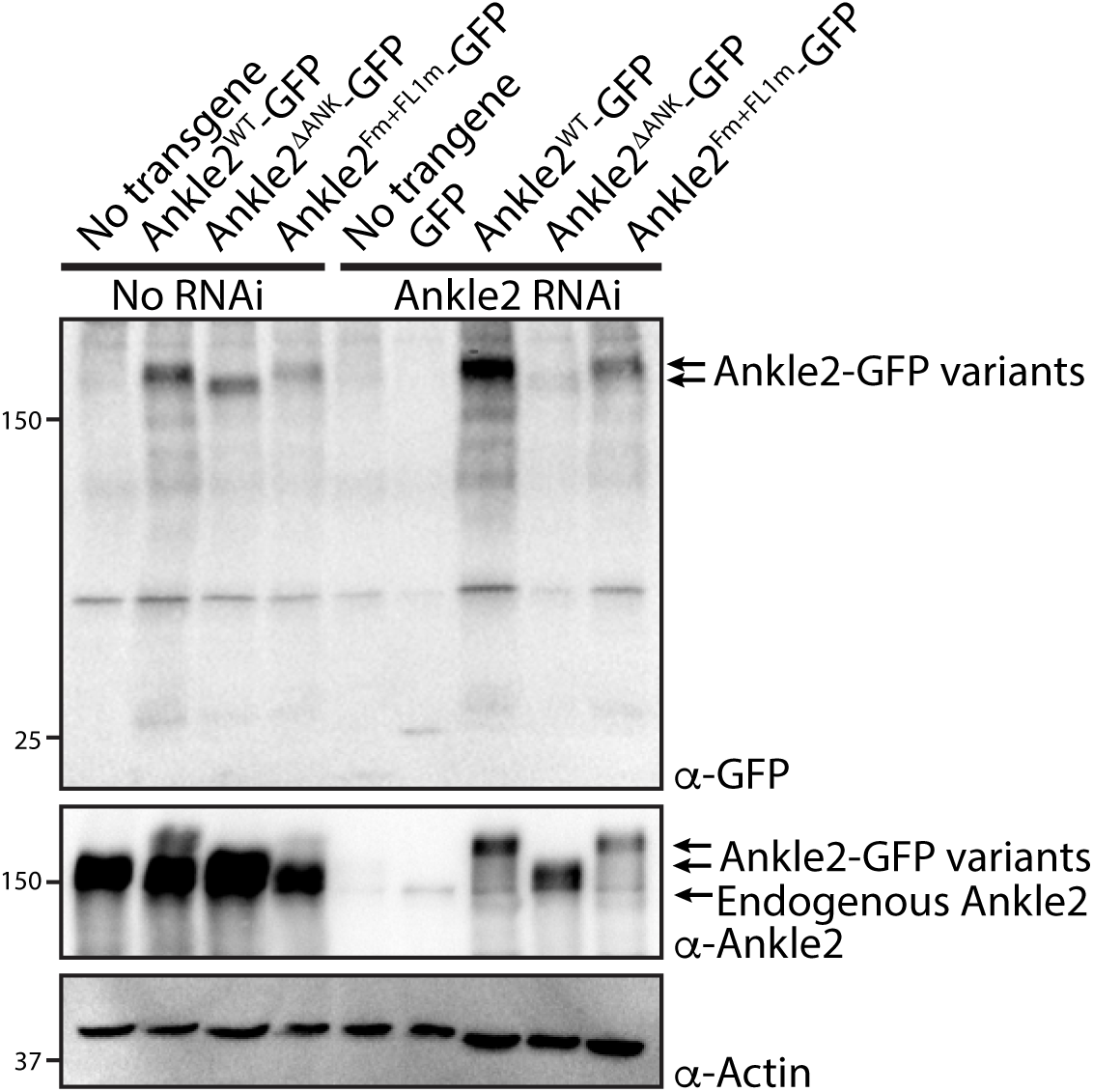
Female germline expression of Ankle2-GFP variants in transgenic lines. Expression of UASp-Ankle2-GFP WT, ΔANK or Fm+FL1m (RNAi insensitive) was induced using the matα4-GAL-VP16 driver. Ankle2 RNAi (line BDSC 77437) was also induced using by the same driver. Western blots from ovaries, probing for GFP, Ankle2 and Actin, show relative expression levels.

**Figure S8.**
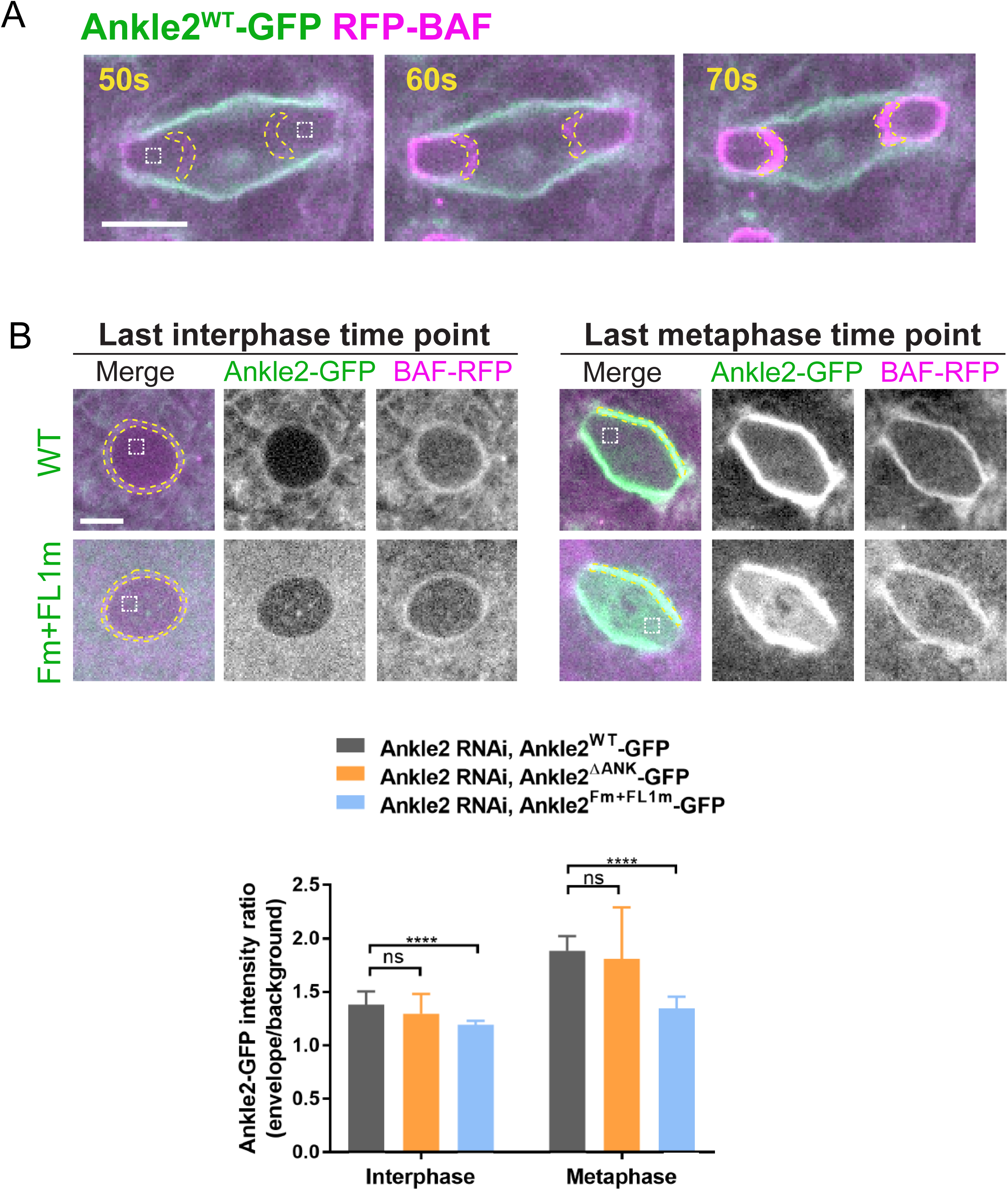
Complement to Fig 6 - Quantifications in embryos. **A.** Illustration of the quantification of fluorescence intensities at the inner core region of the reassembling NE in embryos expressing Ankle2-GFP and RFP-BAF. Inner core regions correspond to the area within the dotted lines where RFP-BAF is recruited in telophase. For each nuclear division quantified, the average of mean intensities from both inner core regions was calculated for each time point. **B.** Interaction of Ankle2 with Vap33 promotes Ankle2 localization during mitosis. Top: Quantification of fluorescence intensities of Ankle2-GFP (WT or Fm+FLm1) at the NE and spindle envelope at the last time points in interphase and metaphase, respectively. Mean intensities of Ankle2-GFP at the NE delineated based on RFP-BAF localization were measured (inside yellow dotted line) in interphase. Fluorescence inside the nucleus was measured as the background (white dotted boxes). For metaphase, mean intensities of Ankle2-GFP in the lateral spindle envelope (inside yellow dotted lines) and within spindles (white dotted boxes, background) were measured. Bottom: ratios of Ankle2-GFP (WT, Fm+FLm1, ΔANK) at the membranes relative to backgrounds in interphase and metaphase. 15 nuclear divisions from 5-6 embryos were quantified for each genotype. All scale bars: 5μm. *** p<0.001, n.s: non-significant (p>0.05) from unpaired t-tests with Welch’s correction.

**Figure S9.**
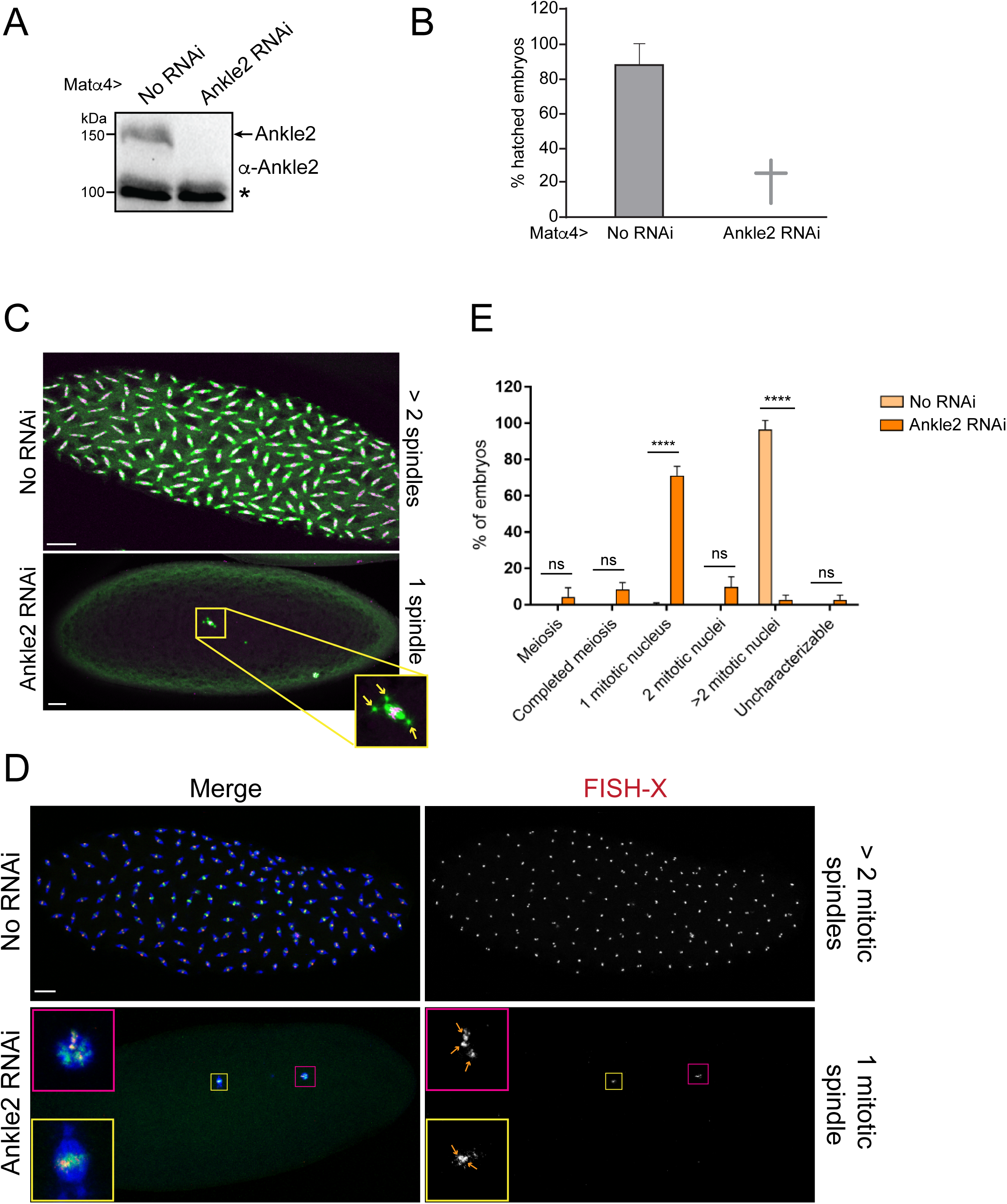
Phenotypes resulting from RNAi depletion of Ankle2 during oogenesis. Ankle2 RNAi (line BDSC77437) was using the matα4-GAL-VP16 driver. **A.** Western blot from ovaries showing Ankle2 depletion. The asterisk indicates a non-specific band used as loading control. **B.** Embryos fail to hatch after Ankle2 RNAi. Averages of 3 independent experiments are shown. Error bar: S.D. **C.** Representative phenotypes of embryos examined by immunofluorescence. Green: α-Tubulin; magenta: histone. Arrows indicate centrosomes, confirming that the spindle is in mitosis and not female meiosis. **D.** Embryos were examined by FISH to reveal the X-chromosome and immunofluorescence for α- Tubulin (blue) and DNA (green). Quantification of embryonic phenotypes. Yellow box: mitotic spindle; pink box: polar body. Enlarged insets are shown on the left. Arrows: X-chromosome foci. **E.** Quantification of the embryonic phenotypes using images as in D. Numbers are averages of 3 independent experiments. At least 50 embryos were scored in each replicate. Error bars: S.D.**** p<0.0001. ns: non-significant from unpaired t-tests. Scale bars: 20 μm.

**Table S1. Proteins identified from affinity purifications of GFP-fused Ankle2 from D-Mel cells.** Numbers of total spectral counts and unique peptides for GFP-Ankle2, Ankle2-GFP or Flag-GFP (control) are indicated.

**Table S2. Proteins identified from affinity purifications of Ankle2-GFP from embryos.** Numbers of total spectral counts and unique peptides for Ankle2-GFP or Flag-GFP (control) are indicated.

**Table S3. Proteins identified from affinity purifications of GFP-fused PP2A-29B from D-Mel cells.** Numbers of total spectral counts and unique peptides for GFP-PP2A- 29B, PP2A-29B-GFP or GFP (control) are indicated.

**Table S4. Proteins identified from affinity purifications of PP2A-29B-GFP from embryos.** Numbers of total spectral counts and unique peptides for PP2A-29B-GFP or GFP (control) are indicated.

**Table S5. Phosphophoptides identification and quantification from cells after Ankle2 or control RNAi.** Results are listed for 4 replicates in each condition, as well as the average fold-changes and p-values.

**Video S1. Localization of Ankle2-RFP and GFP-Vap33 during mitosis and cytokinesis in a D-Mel cell.** Orthogonal maximum intensity projection of 4 z-sections spaced by 1 μm. Images were taken every 205 seconds.

**Video S2. Localization of Ankle2-GFP and RFP-Vap33 during the cell cycle in a syncytial embryo.** A single plane is shown. Images were taken every 10 seconds.

**Video S3. Localization of Ankle2^Fm+FL1m^-GFP and RFP-Vap33 during the cell cycle in a syncytial embryo.** A single plane is shown. Images were taken every 10 seconds.

**Video S4. Localization of Ankle2^WT^-GFP and RFP-BAF during the cell cycle in a syncytial embryo where endogenous is depleted by RNAi.** A single plane is shown. Images were taken every 10 seconds.

**Video S5. Localization of Ankle2^ΔANK^-GFP and RFP-BAF during the cell cycle in a syncytial embryo where endogenous is depleted by RNAi.** A single plane is shown. Images were taken every 10 seconds.

**Video S6. Localization of Ankle2^Fm+FL1m^-GFP and RFP-BAF during the cell cycle in a syncytial embryo where endogenous is depleted by RNAi.** A single plane is shown. Images were taken every 10 seconds.

## REFERENCES

Abramson, J., J. Adler, J. Dunger, R. Evans, T. Green, A. Pritzel, O. Ronneberger, L. Willmore, A.J. Ballard, J. Bambrick, S.W. Bodenstein, D.A. Evans, C.C. Hung, M. O’Neill, D. Reiman, K. Tunyasuvunakool, Z. Wu, A. Zemgulyte, E. Arvaniti, C. Beattie, O. Bertolli, A. Bridgland, A. Cherepanov, M. Congreve, A.I. Cowen-Rivers, A. Cowie, M. Figurnov, F.B. Fuchs, H. Gladman, R. Jain, Y.A. Khan, C.M.R. Low, K. Perlin, A. Potapenko, P. Savy, S. Singh, A. Stecula, A. Thillaisundaram, C. Tong, S. Yakneen, E.D. Zhong, M. Zielinski, A. Zidek, V. Bapst, P. Kohli, M. Jaderberg, D. Hassabis, and J.M. Jumper. 2024. Accurate structure prediction of biomolecular interactions with AlphaFold 3. Nature. 630:493–500.

Alberts, B., A. Johnson, J. Lewis, D. Morgan, M. Raff, K. Roberts, and P. Walter. 2015. Molecular Biology of the Cell. W. W. Norton & Co., United States. 1464 pp.

Archambault, V., J. Li, V. Emond-Fraser, and M. Larouche. 2022. Dephosphorylation in nuclear reassembly after mitosis. Front Cell Dev Biol. 10:1012768.

Asencio, C., I.F. Davidson, R. Santarella-Mellwig, T.B. Ly-Hartig, M. Mall, M.R. Wallenfang, I.W. Mattaj, and M. Gorjanacz. 2012. Coordination of kinase and phosphatase activities by Lem4 enables nuclear envelope reassembly during mitosis. Cell. 150:122–135.

Bobinnec, Y., C. Marcaillou, X. Morin, and A. Debec. 2003. Dynamics of the endoplasmic reticulum during early development of Drosophila melanogaster. Cell Motil Cytoskeleton. 54:217–225.

Deolal, P., J. Scholz, K. Ren, H. Bragulat-Teixidor, and S. Otsuka. 2024. Sculpting nuclear envelope identity from the endoplasmic reticulum during the cell cycle. Nucleus. 15:2299632.

Emond-Fraser, V., M. Larouche, P. Kubiniok, E. Bonneil, J. Li, M. Bourouh, L. Frizzi, P. Thibault, and V. Archambault. 2023. Identification of PP2A-B55 targets uncovers regulation of emerin during nuclear envelope reassembly in Drosophila. Open Biol. 13:230104.

Fishburn, A.T., C.J. Florio, N.J. Lopez, N.L. Link, and P.S. Shah. 2024. Molecular functions of ANKLE2 and its implications in human disease. Dis Model Mech. 17.

Goldberg, M.W. 2017. Nuclear pore complex tethers to the cytoskeleton. Semin Cell Dev Biol. 68:52–58.

Guo, X., X. Dai, X. Wu, T. Zhou, J. Ni, J. Xue, and X. Wang. 2020. Understanding the birth of rupture-prone and irreparable micronuclei. Chromosoma.

Hampoelz, B., and J. Baumbach. 2023. Nuclear envelope assembly and dynamics during development. Semin Cell Dev Biol. 133:96–106.

Haraguchi, T., T. Kojidani, T. Koujin, T. Shimi, H. Osakada, C. Mori, A. Yamamoto, and Y. Hiraoka. 2008. Live cell imaging and electron microscopy reveal dynamic processes of BAF-directed nuclear envelope assembly. J Cell Sci. 121:2540–2554.

Haraguchi, T., T. Koujin, M. Segura-Totten, K.K. Lee, Y. Matsuoka, Y. Yoneda, K.L. Wilson, and Y. Hiraoka. 2001. BAF is required for emerin assembly into the reforming nuclear envelope. J Cell Sci. 114:4575–4585.

Hieda, M. 2019. Signal Transduction across the Nuclear Envelope: Role of the LINC Complex in Bidirectional Signaling. Cells. 8.

Huguet, F., S. Flynn, and P. Vagnarelli. 2019. The Role of Phosphatases in Nuclear Envelope Disassembly and Reassembly and Their Relevance to Pathologies. Cells. 8.

Kaiser, S.E., J.H. Brickner, A.R. Reilein, T.D. Fenn, P. Walter, and A.T. Brunger. 2005. Structural basis of FFAT motif-mediated ER targeting. Structure. 13:1035–1045.

Kamemura, K., and T. Chihara. 2019. Multiple functions of the ER-resident VAP and its extracellular role in neural development and disease. J Biochem. 165:391–400.

Kono, Y., and T. Shimi. 2024. Crosstalk between mitotic reassembly and repair of the nuclear envelope. Nucleus. 15:2352203.

Lancaster, O.M., C.F. Cullen, and H. Ohkura. 2007. NHK-1 phosphorylates BAF to allow karyosome formation in the Drosophila oocyte nucleus. J Cell Biol. 179:817–824.

Li, J., L. Jordana, H. Mehsen, X. Wang, and V. Archambault. 2024. Nuclear reassembly defects after mitosis trigger apoptotic and p53-dependent safeguard mechanisms in Drosophila. PLoS Biol. 22:e3002780.

Li, J., A. Mahajan, and M.D. Tsai. 2006. Ankyrin repeat: a unique motif mediating protein-protein interactions. Biochemistry. 45:15168–15178.

Link, N., H. Chung, A. Jolly, M. Withers, B. Tepe, B.R. Arenkiel, P.S. Shah, N.J. Krogan, H. Aydin, B.B. Geckinli, T. Tos, S. Isikay, B. Tuysuz, G.H. Mochida, A.X. Thomas, R.D. Clark, G.M. Mirzaa, J.R. Lupski, and H.J. Bellen. 2019. Mutations in ANKLE2, a ZIKA Virus Target, Disrupt an Asymmetric Cell Division Pathway in Drosophila Neuroblasts to Cause Microcephaly. Dev Cell. 51:713–729 e716.

Liu, S.Y., and K. Ikegami. 2020. Nuclear lamin phosphorylation: an emerging role in gene regulation and pathogenesis of laminopathies. Nucleus. 11:299–314.

Loewen, C.J., A. Roy, and T.P. Levine. 2003. A conserved ER targeting motif in three families of lipid binding proteins and in Opi1p binds VAP. EMBO J. 22:2025–2035.

Mehsen, H., V. Boudreau, D. Garrido, M. Bourouh, M. Larouche, P.S. Maddox, A. Swan, and V. Archambault. 2018. PP2A-B55 promotes nuclear envelope reformation after mitosis in Drosophila. J Cell Biol. 217:4106–4123.

Murphy, S.E., and T.P. Levine. 2016. VAP, a Versatile Access Point for the Endoplasmic Reticulum: Review and analysis of FFAT-like motifs in the VAPome. Biochim Biophys Acta. 1861:952–961.

Neefjes, J., and B. Cabukusta. 2021. What the VAP: The Expanded VAP Family of Proteins Interacting With FFAT and FFAT-Related Motifs for Interorganellar Contact. Contact (Thousand Oaks). 4:25152564211012246.

Nichols, R.J., M.S. Wiebe, and P. Traktman. 2006. The vaccinia-related kinases phosphorylate the N’ terminus of BAF, regulating its interaction with DNA and its retention in the nucleus. Mol Biol Cell. 17:2451–2464.

Pennetta, G., P.R. Hiesinger, R. Fabian-Fine, I.A. Meinertzhagen, and H.J. Bellen. 2002. Drosophila VAP-33A directs bouton formation at neuromuscular junctions in a dosage-dependent manner. Neuron. 35:291–306.

Petrovic, S., G.W. Mobbs, C.J. Bley, S. Nie, A. Patke, and A. Hoelz. 2022. Structure and Function of the Nuclear Pore Complex. Cold Spring Harb Perspect Biol. 14.

Samwer, M., M.W.G. Schneider, R. Hoefler, P.S. Schmalhorst, J.G. Jude, J. Zuber, and D.W. Gerlich. 2017. DNA Cross-Bridging Shapes a Single Nucleus from a Set of Mitotic Chromosomes. Cell. 170:956–972 e923.

Schellhaus, A.K., P. De Magistris, and W. Antonin. 2016. Nuclear Reformation at the End of Mitosis. J Mol Biol. 428:1962–1985.

Sears, R.M., and K.J. Roux. 2020. Diverse cellular functions of barrier-to-autointegration factor and its roles in disease. J Cell Sci. 133.

Shah, P.S., N. Link, G.M. Jang, P.P. Sharp, T. Zhu, D.L. Swaney, J.R. Johnson, J. Von Dollen, H.R. Ramage, L. Satkamp, B. Newton, R. Huttenhain, M.J. Petit, T. Baum, A. Everitt, O. Laufman, M. Tassetto, M. Shales, E. Stevenson, G.N. Iglesias, L. Shokat, S. Tripathi, V. Balasubramaniam, L.G. Webb, S. Aguirre, A.J. Willsey, A. Garcia-Sastre, K.S. Pollard, S. Cherry, A.V. Gamarnik, I. Marazzi, J. Taunton, A. Fernandez-Sesma, H.J. Bellen, R. Andino, and N.J. Krogan. 2018. Comparative Flavivirus-Host Protein Interaction Mapping Reveals Mechanisms of Dengue and Zika Virus Pathogenesis. Cell. 175:1931–1945 e1918.

Shevelyov, Y.Y. 2023. Interactions of Chromatin with the Nuclear Lamina and Nuclear Pore Complexes. Int J Mol Sci. 24.

Slee, J.A., and T.P. Levine. 2019. Systematic prediction of FFAT motifs across eukaryote proteomes identifies nucleolar and eisosome proteins with the predicted capacity to form bridges to the endoplasmic reticulum. Contact (Thousand Oaks). 2:1–21.

Snyers, L., R. Erhart, S. Laffer, O. Pusch, K. Weipoltshammer, and C. Schofer. 2018. LEM4/ANKLE-2 deficiency impairs post-mitotic re-localization of BAF, LAP2alpha and LaminA to the nucleus, causes nuclear envelope instability in telophase and leads to hyperploidy in HeLa cells. Eur J Cell Biol. 97:63–74.

Thomas, A.X., N. Link, L.A. Robak, G. Demmler-Harrison, E.C. Pao, A.E. Squire, S. Michels, J.S. Cohen, A. Comi, P. Prontera, A. Verrotti di Pianella, G. Di Cara, L. Garavelli, S.G. Caraffi, C. Fusco, R. Zuntini, K.C. Parks, E.H. Sherr, M.O. Hashem, S. Maddirevula, F.S. Alkuraya, I.A.F. Contractar, J.E. Neil, C.A. Walsh, H.J. Bellen, H.T. Chao, R.D. Clark, and G.M. Mirzaa. 2022. ANKLE2-related microcephaly: A variable microcephaly syndrome resembling Zika infection. Ann Clin Transl Neurol. 9:1276–1288.

Ungricht, R., and U. Kutay. 2017. Mechanisms and functions of nuclear envelope remodelling. Nat Rev Mol Cell Biol. 18:229–245.

Velez-Aguilera, G., S. Nkombo Nkoula, B. Ossareh-Nazari, J. Link, D. Paouneskou, L. Van Hove, N. Joly, N. Tavernier, J.M. Verbavatz, V. Jantsch, and L. Pintard. 2020. PLK-1 promotes the merger of the parental genome into a single nucleus by triggering lamina disassembly. Elife. 9.

Xu, Y., Y. Chen, P. Zhang, P.D. Jeffrey, and Y. Shi. 2008. Structure of a protein phosphatase 2A holoenzyme: insights into B55-mediated Tau dephosphorylation. Mol Cell. 31:873–885.

Yamamoto, S., M. Jaiswal, W.L. Charng, T. Gambin, E. Karaca, G. Mirzaa, W. Wiszniewski, H. Sandoval, N.A. Haelterman, B. Xiong, K. Zhang, V. Bayat, G. David, T. Li, K. Chen, U. Gala, T. Harel, D. Pehlivan, S. Penney, L. Vissers, J. de Ligt, S.N. Jhangiani, Y. Xie, S.H. Tsang, Y. Parman, M. Sivaci, E. Battaloglu, D. Muzny, Y.W. Wan, Z. Liu, A.T. Lin-Moore, R.D. Clark, C.J. Curry, N. Link, K.L. Schulze, E. Boerwinkle, W.B. Dobyns, R. Allikmets, R.A. Gibbs, R. Chen, J.R. Lupski, M.F. Wangler, and H.J. Bellen. 2014. A drosophila genetic resource of mutants to study mechanisms underlying human genetic diseases. Cell. 159:200–214.

